# Dynein synergises with EB1 to facilitate cortex-microtubule encounter and proper spindle positioning in metaphase

**DOI:** 10.1101/2025.05.12.653564

**Authors:** Christoforos Efstathiou, Nikola Ojkic, Viji M. Draviam

**Affiliations:** Centre for Cell Dynamics, School of Biological and Behavioural Sciences, Queen Mary University of London, London E1 4NS; London Interdisciplinary Doctoral Training Program

## Abstract

To rotate the bulky mitotic spindle, astral microtubules are pulled by cortex-bound dynein. We investigate whether dynein regulates astral microtubules, independent of its classical microtubule-pulling role by combining super-resolution live-cell microscopy with optogenetic tools to disrupt and measure astral microtubule distributions and spindle movements. Localised photoinactivation of EB1 reveals that microtubule ends reaching distinct cortical regions differently influence spindle movements. Using a dynein inhibitor gradient, we separate dynein’s role in spindle positioning (maintaining pole-to-cortex distance) from its role in directing spindle movements. EB1 photoinactivation of whole spindles uncovers a previously unrecognised synergistic role for dynein and EB1 in maintaining astral microtubules reaching the cortex. We demonstrate the EB1-dependent role for dynein in maintaining astral microtubule growth is independent of its classical cortical roles by analysing dynamic changes in spindle position, displacement rates and cortical dynein localisation and by depleting dynein’s cortical platform, LGN. Our finding that Dynein synergises with EB1 to regulate astral microtubule, independent of cortical pulling, has implications for how spindle positioning is correctly established in cells.

**HIGHLIGHTS:**

- Systematic quantification of spindle movements and microtubule lengths reveals dynein’s dual role in microtubule growth and pulling.
- Microtubules reaching distinct cortex regions differently regulate spindle movements.
- EB1 photoinactivation shows dynein promotes EB1-independent astral microtubule growth.
- Reduced dynein causes shorter asters and asymmetric spindle positioning, without reducing spindle movements.
- Noncortical dynein regulates cortex-microtubule encounter; cortical dynein controls spindle movement.

## INTRODUCTION

The final position of the spindle defines the plane of cell division. Spindle positioning errors can cause daughter cells of irregular sizes or in incorrect positions within tissues[1–4]. Spindle positioning is determined by a cortex-based force-generation machinery, including the minusend directed motor dynein, and its partner dynactin which are enriched at the cell cortex by the NuMA-LGN-G*α*_i_ complex [5–12]. Although small molecule inhibitors of dynein are available [13, 14], they have not been exploited to dissect dynein’s roles in spindle movement and positioning. Spindle positions can change in response to cell shape [9, 15], mechanical stress [16], morphogenesis cues [17] and tissue tension [18]. How these signals converge on dynein’s function to guide spindle movements is not fully understood.

Biophysical models include two factors that control spindle movements: microtubule dynamics and cortical-cue interactions [16, 19–22]. In agreement with the models, the cortical polarity kinase Par1 (MARK2) orients the spindle in 3Dimensions (3D) by centering the spindle through microtubule growth, independent of cortical dynein [23], and aligning along the substratum by clustering dynein [21] along the cortical midzone [24]). Apart from cortex localisation, Dynein-Dynactin is recruited to microtubule-ends by EB1 and EB3 [25, 26]. While dynein-coated substrates pull at microtubules [7, 27, 28], microtubules promote dynein’s cortical recruitment[29], reinforcing each other, hindering the separation of dynein’s mitotic roles in microtubule pulling, tethering, and growth.

EB proteins [30, 31] share three domains, N-terminal CH domain [32, 33] to bind microtubules, EB-homology domain to bind proteins possessing SxIP motif [34–38] and C-terminal EEY-motif [39–41] to bind CAP-Gly proteins. Yet they have non-overlapping cellular roles [25, 33, 41–50]. Considering the distinct location of EB3 trailing behind EB1 at microtubule-ends [48], acute and reversible photoinactivation of EB1 [50, 51] allows the probing of functional differences between EB3 and EB1 in astral microtubule regulation. Combining transient EB1 photoinactivation and small-molecule dynein inhibitor gradients can facilitate systematic combinatorial analysis of microtubule regulation and motor function in orchestrating spindle positions and movements, and this approach has not been explored so far.

Here we report a role for dynein in astral microtubule length regulation independent of its well-established role in microtubule pulling. Using super-resolution live-cell microscopy, we show that astral microtubules retrace paths to encounter cortical interaction sites. Localised photoinactivation of growing microtubules that reach different areas of the cortex shows their differing roles in the position and movement of mitotic spindles. EB1 photoinactivation shows that EB3 cannot fully compensate for the lack of EB1, and EB1 synergises with dynein in maintaining astral microtubules: partial loss of dynein activity accelerates microtubule shrinkage in monopolar spindles. This role of dynein can be recapitulated in bipolar spindles, since partial loss of dynein reduces pole-to-cortex distances causing shorter asters and asymmetrically positioned spindles, without altering cortex-bound dynein levels. In agreement, in LGN-depleted cells lacking cortical dynein, we find no significant reduction in astral microtubule length in the presence or absence of EB1. Together, our studies reveal a role for non-cortical dynein in facilitating astral microtubule interaction at the cell cortex.

## RESULTS

### Distinct roles for microtubules reaching different cortical regions

EB3-decorated microtubule-ends interact with the cortex [52, 53], but microtubule-ends of the bipolar spindle are highly dynamic and crowded making their tracking challenging[54]. To overcome this, we used monopolar spindles that display astral microtubules extending throughout the cortex. To study cortex-microtubule interactions in monopolar spindles, we used Structured Illumination Microscopy (SIM^2^) that can resolve up to 60 nm resolution. In a H1299 cell line expressing a blue-light sensitive EB1 variant (*π*-EB1) [51, 55, 56], we added an EB3-mKate2 marker (H1299 *π*-EB1-EGFP/EB3-mKate2) to observe growing microtubule-ends. Super-resolved movies show EB3 comets followed similar paths towards the cortex, occasionally converging at the same cortical site (n=16 bipolar and 10 monopolar spindles, Figure S1 A). This observation is consistent with cortical dynein-mediated astral microtubule capture at specific cortical sites [21, 29, 57]. EB3 comets that reached the cortex show a change in morphology, with smaller comets retracing the path (Figure S1A) in both monopolar and bipolar spindles.

To understand the link between cortex-microtubule interactions and spindle movements, we photoinactivated EB1 [51, 55] in our H1299 *π*-EB1-EGFP/EB3-mKate2 cell line. We first set up whole-cell EB1 photoinactivation using blue light illumination and image-acquisition blocks (Figure S1B). In both bipolar and monopolar spindles, following EB1 inactivation, EB3-mKate2 comets continued to localise at the microtubule-end, although more diffuse compared to the Dark state (without blue light) (Figure S1 A), allowing us to track microtubule plus-ends. Following 200s of EB1 photoinactivation, the bipolar spindle size (pole-to-pole distance) was reduced confirming that EB3 can not fully compensate for the lack of EB1 in maintaining bipolar spindle length, as reported [51]. In monopolar spindles, EB1 photoinactivation resulted in a shrinking monopolar spindle (Figure S1 B). These suggest non-redundant roles for EB1 and EB3 in both monopolar and bipolar spindles.

To selectively perturb cortex-microtubule interactions, we set up a ‘localised’ blue light illumination system by coupling the super-resolution SIM system with an independent photokinetics laser module for spatially restricted illumintation [58] (Figure 1A). The laser intensity was optimised to minimally affect the EB3-mKate2 intensities during EB1 photoinactivation. To validate the setup, we used H1299 mCherry-Zdk1-EB1C cells [51] to visualise non-photoinactivated EB1 at (561 nm) and photoinactivate a specified Region Of Interest (ROI) with a 473 nm laser (Figure 1A). Our setup could locally photoinactivate EB1 in flat interphase and rounded-up mitotic cells (Figure 1A). In bipolar spindles, exposing only one spindle pole to blue light, and asymmetrically photoinactivating EB1, resulted in the shrinkage of EB3 signals at the illuminated pole, which aligns with previous observations [51] (Figure 1A), confirming the efficacy of the super-resolution set up. We photoinactivated interphase cell periphery in H1299 *π*-EB1EGFP/EB3-mKate2 (Figure 1B). EB3 comets continued to grow into the blue light-illuminated region where EB1 was acutely photoinactivated (Figure 1B). However, the length of the EB3 comets appeared to steadily decrease upon entering the blue light region (Figure 1B), similar to whole-cell studies (Figure S1 B) and previous reports [55]. Localised EB1 photoinactivation with EB3 tracking allowed us to observe microtubule-ends with a reduction in EB3 signal. Thus, EB3 can localise and maintain the growth of microtubule-ends independent of EB1, despite a visible impact on EB3 comet length, suggesting a reduction in microtubule polymerisation.

**Figure 1.**
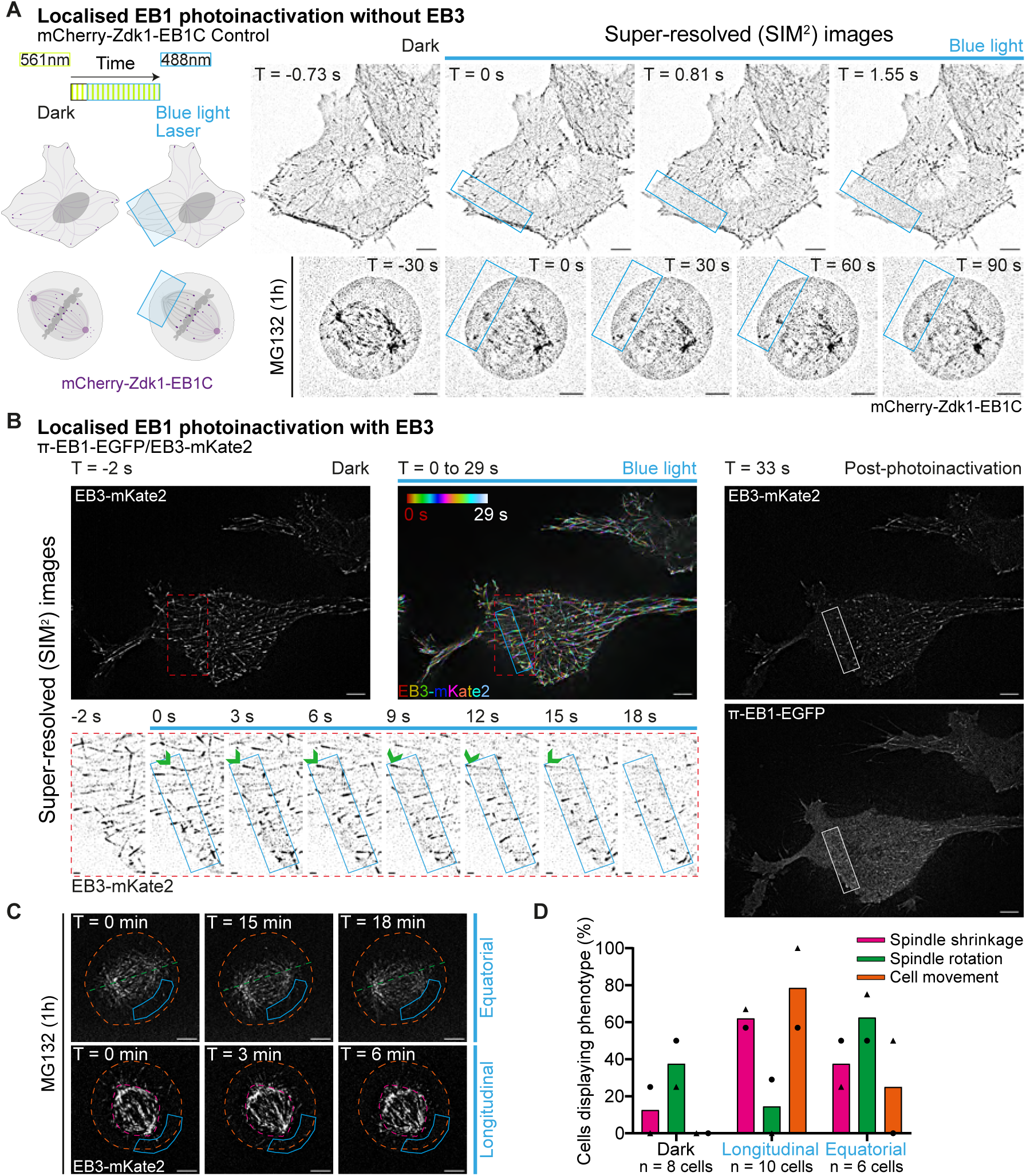
EB3 and EB1 operate independently with distinct roles for microtubules reaching equatorial and polar cortical regions. **A.** Schematic illustrating localised EB1 photoinactivation (blue rectangle) via a photokinetics laser module. Next to the schematic are representative lattice SIM^2^ single slice time-lapse images of H1299 mCherry-Zdk1-EB1C interphase and mitotic cell undergoing localised EB1 photoinactivation (blue rectangle) at T = 0 s. Images are representative of 26 cells across two experimental repeats. **B.** Representative lattice SIM^2^ time-lapse images of an H1299 *π*-EB1-EGFP/EB3-mKate2 cell in interphase cell exposed to blue light locally (blue rectangles) using a photokinetics laser module. Temporally colour coded image maximum projection image shows localised EB1 photoinactivation (blue box). Within the red dotted box are magnified time-lapse image sequences (inverted contrast) of the region indicated, showing EB3 microtubule ends (green arrow) growing through the blue light-exposed area (blue rectangles). The white rectangle indicates the blue light-exposed area post-photoinactivation. Images are representative of 14 cells across two experimental repeats. **C.** Representative lattice SIM^2^ time-lapse images of H1299 *π*-EB1EGFP/EB3-mKate2 bipolar spindles undergoing localised longitudinal (behind spindle pole) or equatorial (below the spindle pole-pole axis) blue light exposure (dotted blue polygon) via a photokinetics laser module. Cells were treated with MG132 1 h before imaging, and DMSO or control siRNA just before imaging. Green dotted line and orange dotted circle show initial (T = 0 min) pole-pole axis orientation and cell position, respectively. Magenta dotted polygon shows original (T = 0 min) spindle shape. **D.** Bar chart quantifying the mean proportion of spindle and cell behaviour without EB1 photoinactivation (Dark, as control), and with localised longitudinal and equatorial EB1 photoinactivation in H1299 *π*-EB1-EGFP/EB3-mKate2 bipolar spindles. Different shaped points represent different experimental repeats. Sample size (n) as indicated across two experimental repeats. T corresponds to the time at which the image was taken. Scale bar = 5 µm, 1 µm for zoomed images.

Next we compared the roles of microtubule-ends reaching different cortical regions. In metaphase-arrested H1299 *π*-EB1-EGFP/EB3-mKate2 cells, we photoinactivated using two patterns: either behind one of the poles (longitudinal) or near the equator of the spindle (equatorial) (Figure 1C). Spindle tumbling was not observed in localised EB1 photoinactivation studies regardless of photoinactivation in the polar region or not (n=16 cells). EB1 photoinactivation in the polar region (longitudinal) primarily resulted in spindle shrinkage (n=10 cells) (Figure 1C and D). Conversely, EB1 photoinactivation in the equatorial region resulted in a spindle rotational movement (Figure 1C and D). These findings show that microtubules reaching the equatorial cortex can impact spindle movements, and microtubules reaching polar cortex are exquisitely sensitive to the loss of EB1. These suggest a separation of spindle movement and microtubule regulation, based on the cortical region.

### Partial loss of dynein reduces spindle pole-to-cortex distances, without reducing spindle displacement or altering cortical dynein localisation

To separate dynein’s roles in spindle maintenance and movements, we exploited dynein inhibitor gradients. MG132 treated metaphase HeLa cells expressing DHC-GFP [24] were exposed to two different dynein inhibitors, Dynarrestin (Dyn) [13] and Ciliobrevin D (CilioDi) [14], at three different concentrations (Figure S2 A). Tracking spindle poles relative to the cortex using DHC-GFP as a marker showed the characteristic oscillatory movement of spindles along the pole-pole axis [9, 23, 59]. We binned spindle movements into three levels: ‘substantial’ with more than 2 spindle oscillations; ‘limited’, with at least 1 spindle oscillation; and ‘None’, without any movement in the 60 min movie (Figure S2B and C). Cortical DHC-GFP was not affected by Ciliobrevin, as previously shown [57], or Dynarrestin treatments (Figure S2A). Taking advantage of cortical DHC-GFP localisation, we additionally assessed whether DHC-GFP localisation is ‘dynamic’ (shifting from one side of the cortex to the other) or ‘fixed’ (remained on one side of the cortex) (1 h) (Figure S2A and C).

Low Ciliobrevin-D, in the presence of serum, allowed spindle movement similar to controls. At least 50 % of cells (Ciliobrevin-D 5 µM 52 %, Ciliobrevin-D 500 nM 65 %, Ciliobrevin-D 50 nM 63 % vs DMSO 54 %) exhibited more than 2 spindle oscillations within 60 min (substantial; Figure S2A and S2B). The spindle oscillatory movement after low-levels of Ciliobrevin-D-treatment was reflected in DHC-GFP localisation, as more than 80 % of cells (Ciliobrevin-D 5 µM 92 %, Ciliobrevin-D 500 nM 92 %, Ciliobrevin-D 50 nM 84 % versus DMSO 89 %) showed dynamic cortical DHC-GFP crescent localisation, correlating with spindle pole positions (Figure S2A and S2C). These are consistent with studies showing that 10 µM to 75 µM Ciliobrevin-D abrogates spindle movements [16, 57], and 10 µM Ciliobrevin-D partially inhibits dynein without disrupting spindle bipolarity [60].

High concentrations of Dynarrestin (2.5 µM and 25 µM) significantly (p < 0.05 limited, p < 0.01 substantial movement vs DMSO) impaired spindle movement, where more than 65 % of cells (Dynarrestin 25 µM n = 28 cells, Dynarrestin 2.5 µM n = 24 cells) showed no oscillations (Figure S2B). 2.5 µM or 25 µM Dynarrestin-treated cells also showed limited movement (Dynarrestin 25 µM 35 %, Dynarrestin 2.5 µM 16 %). Thus, 2.5 µM of Dynarrestin can also inhibit dynein-mediated spindle pulling without disrupting spindle morphology.

Following low-levels of 250 nM Dynarrestin treatment, 48 % of cells showed limited movement (at least 1 spindle oscillation), while 28 % displayed substantial movement (vs DMSO 54 %) (Figure S2B), indicating that 250 nM Dynarrestin inhibits dynein partially, allowing cortical dynein-mediated spindle pulling. As expected, spindle movements observed at different Dynarrestin concentrations affected DHC-GFP crescent dynamics, with 2.5 µM and 25 µMtreated cells exhibiting no shift across the opposing cortices (Dynarrestin 25 µM 85 %, Dynarrestin 2.5 µM 87 % vs DMSO 11 %; p < 0.001), relative to DMSO (Figure S2A and C). Thus, Ciliobrevin-D and Dynarrestin concentration gradients allow the dissection of dynein controlled spindle positions and movement.

Unlike DHC-GFP, EB3-mKate2 allows the use of our AI-guided spindle segmenting software SpinX [61] to precisely track spindle movements in 3D (Figure S3A). Quantifying pole-to-cortex (pole-cortex) distances revealed a significant decrease after Ciliobrevin-D 50 nM treatment (DMSO 3.28 µm vs Ciliobrevin-D 50 nM 2.99 µm; median) (Figure S3D), with no change in spindle size (Figure S3A), leading to a short aster and asymmetrically positioned spindle. Importantly, there was no significant difference in total spindle displacement for a period of 30 min between DMSO and Ciliobrevin-D 50 nM-treated cells (DMSO 65.4±20.7 µm vs CiliobrevinD 50 nM 65.9±22.1 µm; mean±SD) (Figure S3B). In addition, there was no significant difference in cumulative spindle displacement between DMSO and Ciliobrevin-D 50 nM treatments (DMSO vs Ciliobrevin-D 50 nM p = 0.96; Mann-Whitney U test) (Figure S3C). To ascertain if asymmetric spindle positioning (differences in pole-cortex distances) arose from differences in spindle pulling towards the opposing cortex, we used SpinX tracker output to measure spindle exit velocities and reflection angles when a pole closest to the cortex moved away from the cortex (Figure S3E). Velocities after reflection and reflection angles (direction of movement) are both similar in control and ciliobrevin D treated cells (DMSO: 1.3+/-0.8 micron/min, Ciliobrevin-D 1.2+/-0.7 micron/min) (Figure S3F and S3G), suggesting no significant changes in dynein-mediated pulling or steering of spindle movements.

To correlate spindle movement, spindle pole position, cortical dynein status and dynein activity using Hela DHC-GFP cells were exposed to the lowest concentrations of two dynein inhibitors, Ciliobrevin-D 50 nM and Dynarrestin 250 nM. Spindle movements were assessed by manual tracking of spindle poles using time-lapse movies of cells treated with MG132 as before [24] (Figure 2A). Spindles displaying less than 1 oscillation (within 60 min) along the polepole axis, in Dynarrestin 250 nM-treated cells were excluded from the analysis. No significant differences were observed in instantaneous displacement of spindles in control and dynein inhibitor-treated cells (DMSO 3.67±2.08 µm vs Ciliobrevin-D 50 nM 4.17±2.63 µm, DMSO 4.13±2.62 µm vs Dynarrestin 250 nM 4.32±2.87 µm; mean±SD) (Figure 2B and 2C). No significant differences were observed in the total or cumulative displacement of spindles in control and dynein inhibitor-treated cells (DMSO 73.4±17.6 µm vs Ciliobrevin-D 50 nM 83.4±25.6 µm, DMSO 86.3±23.3 µm vs Dynarrestin 250 nM 82.6±21.1 µm; mean±SD); cells exposed to Ciliobrevin-D 50 nM showed a slight cumulative increase (1.14 times) and cells exposed to Dynarrestin 250 nM showed a marginal decrease (0.95 times) (Figure 2D). Similarly, comparing the distribution of individual spindle pole displacement rates (over sequential time points) showed no significant differences in the peak values or distribution of rates (peaks of kernel density estimation (KDE) curves) (Figure 2E), confirming no reduction in spindle displacement at low concentrations of Dynein inhibitor treatments.

**Figure 2.**
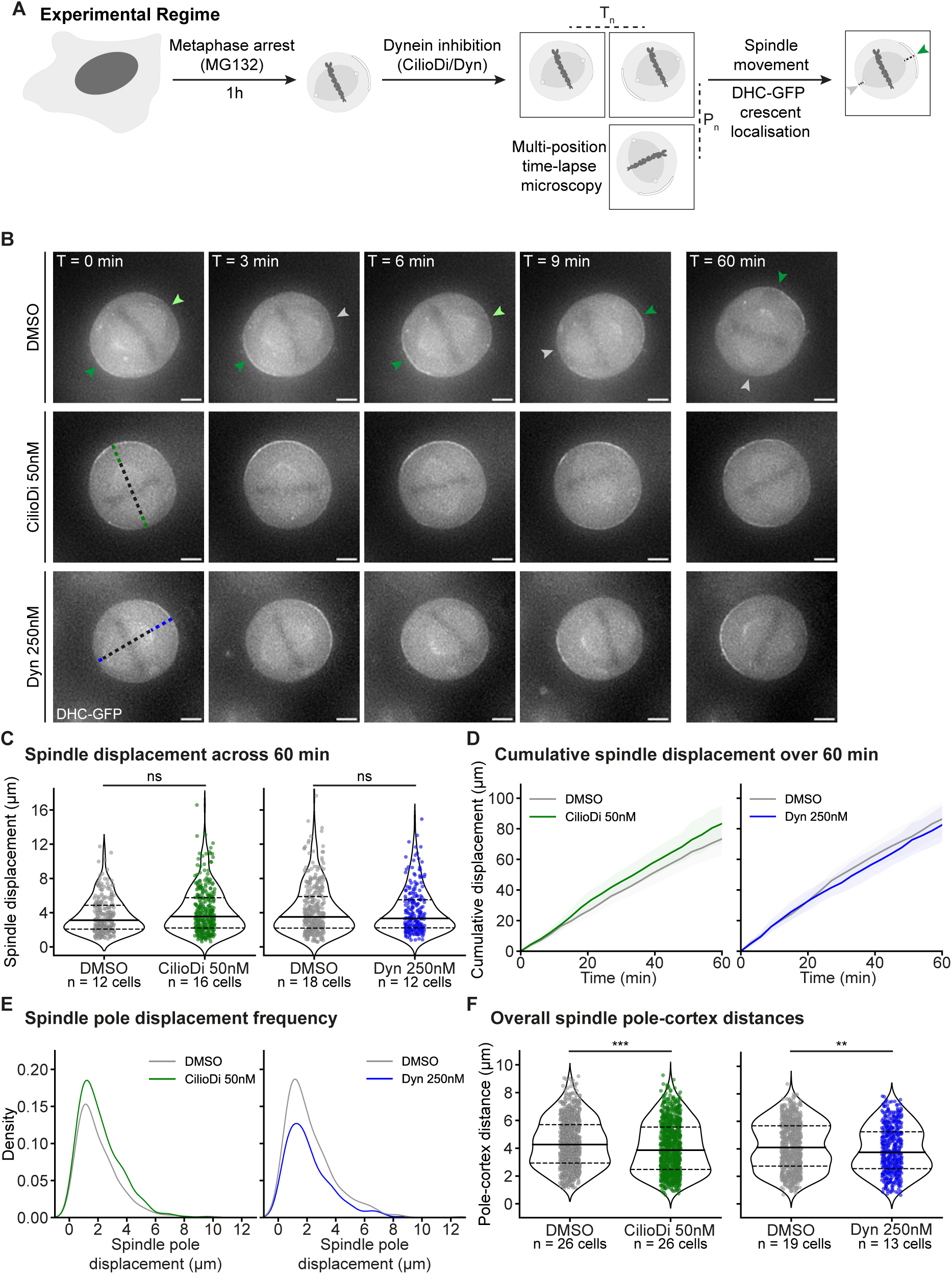
Partial inhibition of dynein reduces spindle pole-cortex distances without altering total displacement. **A.** Experimental regime. HeLa DHC-GFP cells were treated with MG132 1 h before imaging, and either DMSO (Control) or Ciliobrevin D (Ciliobrevin-D 5 µM, 500 nM, 50 nM) or Dynarrestin (Dynarrestin 25 µM, 2.5 µM, 250 nM) just before imaging. Multiple positions (P_n_) were imaged across wells at 3 min intervals for a total time of 60 min (T_n_). Cells were filmed for up to 3 h. Overall spindle pole-to-pole movement and DHC-GFP cortical crescent localisation status were visually assessed. Green arrowhead signifies DHC-GFP crescent localisation, while grey arrowhead indicates no DHC-GFP crescent localisation. **B.** Representative widefield deconvolved maximum projected time-lapse images of HeLa DHC-GFP cells treated with MG132 and either DMSO (Control) or Ciliobrevin-D 50 nM or Dynarrestin 250 nM. Green arrowhead marks cortical DHC-GFP crescent localisation, light green arrowhead shows faint cortical DHC-GFP localisation, and grey arrowhead indicates no prominent cortical DHC-GFP crescent localisation. Dotted black lines indicate the spindle pole-pole axis, while green and blue dotted lines display spindle pole-cortex distances for Ciliobrevin-D 50 nM or Dynarrestin 250 nM respectively. **C.** Violin plots showing the distribution of instantaneous displacement exhibited by spindles treated with DMSO, Ciliobrevin-D 50 nM and Dynarrestin 250 nM over a 60 min imaging period. DMSO vs Ciliobrevin-D 50 nM consists of 2 experimental repeats, while DMSO vs Dynarrestin 250 nM consists of 3. Dots indicate individual measurements. **D.** Line plots quantifying the average cumulative displacement exhibited by spindles treated with DMSO, Ciliobrevin-D 50 nM and Dynarrestin 250 nM. Shaded region displays 95 % confidence interval. **E.** KDE curves showing the density distribution of spindle pole displacement values over sequential time points for spindles treated with DMSO, Ciliobrevin-D 50 nM and Dynarrestin 250 nM. Only displacement values of the pole farthest from the cortex at the initial time point were included. **F.** Violin plots showing the distribution of pole-cortex distances in DMSO, Ciliobrevin-D 50 nM and Dynarrestin 250 nM-treated spindles. Dots represent measurements from both spindle poles across all timepoints. Dotted lines indicate the quartiles, while the solid lines show the median. Data for both comparisons are across 3 experimental repeats. In (**C**) no statistical significance (ns) was determined by unpaired t-test after confirming equal variances with Levene’s test. No statistical significance in (**D**) and statistical significance (*) in (**E**) was determined through a Mann-Whitney U test. Normality was checked with a Shapiro-Wilk test. T corresponds to the time at which the image was acquired. Scale bar = 5 µm. Sample size (n) displayed in (**C**) is the same for (**D**). Sample size in (**E**) was DMSO 20 cells/319 values vs Ciliobrevin-D 50 nM 26 cells/416 values; DMSO 19 cells/371 values vs Dynarrestin 250 nM 13 cells/256 values. Sample size in (**F**) as shown.

In stark contrast, cells treated with Ciliobrevin-D 50 nM and Dynarrestin 250 nM exhibited statistically significant reduction in pole-cortex distances compared to the control (DMSO vs. Ciliobrevin-D 50 nM p = 0.0002, DMSO vs. Dynarrestin 250 nM p = 0.0053; Mann-Whitney U test) (Figure 2F). Despite a modest magnitude of 7 percentage reduction in pole-to-cortex distance ((DMSO 4.38±1.75 µm vs Ciliobrevin-D 50 nM 4.06±1.83 µm, DMSO 4.21±1.74 µm vs Dynarrestin 250 nM 3.93±1.69 µm; mean±SD), this consistent and statistically significant difference is important, as it translates to a 19.6 percentage reduction in the volume of the spherical cap (relevant to the search space for astral microtubule-ends encountering the cortex). It would also mean a relative asymmetry of 39.2 percentage between the search space for astral microtubule-ends emnating from the two spindle poles. Reduction in pole-to-cortex distance following partial reduction in dynein activity are consistent with our observations in the EB3-mKate2 cell line (Figure S3D). Thus, using different inhibitors and different cell lines, we show a role for dynein in maintaining pole-cortex distances (Figure 2F). In summary, asymmetric spindle positioning and reduced pole-to-cortex distance observed following partial loss of dynein is independent of previously reported roles of dynein in spindle pulling and displacement.

To understand the reason for reduced pole-cortex distance, we assessed specifically the movement of the pole closest to the cortex, and found a significant reduction in pole-cortex distances upon Ciliobrevin-D 50 nM and Dynarrestin 250 nM treatment (Figure S4A). This was intriguing since we expected a reduction in cortical pull following dynein inhibition which would position the pole away from the cortex following dynein inhibition. So, we analysed the relationship between spindle positions and cortical DHC-GFP localisation, by grouping pole-cortex distances according to the DHC-GFP crescent as ‘Crescent’, ‘Faint’, or ‘No Crescent’ (Figure S4B). In both inhibitor treatments, there was a significant reduction in pole-cortex distances when DHC-GFP crescent was present (Crescent: DMSO 5.26 µm vs Ciliobrevin-D 50 nM 4.91 µm, DMSO 5.26 µm vs Dynarrestin 250 nM 4.88 µm; median), as well as absent (No Crescent: DMSO 3.25 µm vs Ciliobrevin-D 50 nM 2.62 µm, DMSO 3.11 µm vs Dynarrestin 250 nM 2.68 µm; median) (Figure S4B). These show that the role of dynein in maintaining pole-cortex distances can be independent of its cortical localisation. Nevertheless, cortical DHCGFP status is dependent on pole position, consistent with the current paradigm [59, 62–64]: as the pole moves closer to the cortex displaying DHC-GFP enrichment, DHC-GFP signals begin to reduce (i.e. ‘No Crescent’) (Figure 2B and Figure S4C). Prior reports of pole-cortex distances and cortical DHC-GFP enrichment observed DHC-GFP crescent when the pole was typically 3.6 µm away from the cortex, with a GFP signal loss occurring by 1.4 µm [59]. Here DHC-GFP crescent enrichment was maximal at a distance of 5.34 µm (Crescent: DMSO left plot 5.08 µm and DMSO right plot 5.60 µm; peaks of KDE curves), while signal dissipation occurred around 2.97 µm (No Crescent: DMSO left plot 2.86 µm and DMSO right plot 3.08 µm; peaks of KDE curves) (Figure S4C). Although the pole-cortex distances observed are greater than the reported values [59], the difference in distance between the presence or absence of DHC-GFP crescent is similar (2.2 µm [59] vs 2.37 µm, this study).

Spindle poles associated with no DHC-GFP crescent displayed a peak pole-cortex distance that was lesser in dynein inhibitor treated cells compared to controls (No Crescent: DMSO 2.86 µm vs CilioDi 50 nM 2.27 µm, DMSO 3.08 µm vs Dyn 250 nM 2.54 µm; peaks of KDE curves) (Figure S4C), confirming that poles spent a longer period closer to the cortex following partial loss of dynein. Collectively partial loss of dynein reduced pole-cortex distances without altering total spindle displacement, and cortical dynein localisation.

### Dynein and EB1 facilitate astral microtubule length maintenance

To investigate whether reduced pole-cortex distance was due to reduced astral microtubules, we analysed the efficiency with which astral microtubules reached the cortex following partial loss of dynein, in the presence and absence of EB1. To explore this, quantitatively, we tracked astral microtubule growth in monopolar spindles following 4 h STLC treatment [24] in H1299 *π*EB1-EGFP/EB3-mKate2 cells (Figure 3A). Midplane images of the monopolar spindle captured at frequent time intervals (4 s) allowed the tracking of microtubule-end decorated EB3 comets while minimising phototoxicity and photobleaching over an extended period (10 min, details in Methods). To quantify EB3-decorated microtubule-ends encountering the cortex, we optimised a workflow using TrackMate’s LoG detector [65] which was robust in detecting the furthest-reaching comets across time (Figure S5A-C). We first compared the monopolar spindle area either without blue light or only in the first frame before blue light exposure (dark state) (Figure S5D). The ratio of monopolar spindle area to cell area did not show any significant difference between control and dynein inhibition conditions (DMSO 0.66±0.09 vs Ciliobrevin-D 50 µM 0.69±0.13, Ciliobrevin-D 5 µM 0.62±0.08, Dynarrestin 25 µM 0.64±0.15, Dynarrestin 2.5 µM 0.56±0.15; mean±SD ratio) (Figure S5D), suggesting that microtubule ends can grow through the cytoplasm to reach the cortex regardless of reduction in dynein activity.

**Figure 3.**
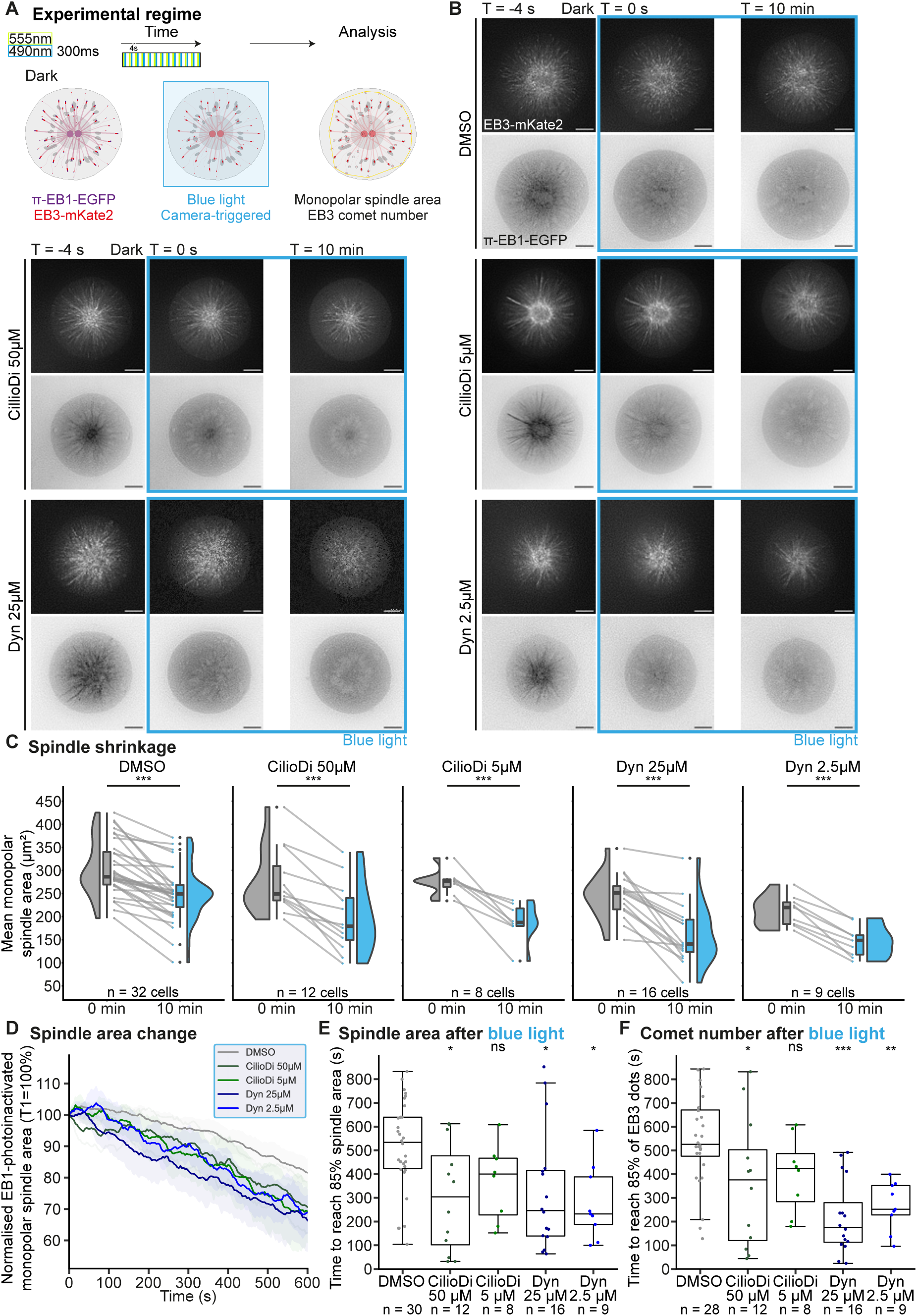
Dynein and EB1 maintain astral microtubule length. **A.** Schematic illustrating camera-triggered whole cell EB1 photoinactivation (blue border) in H1299 *π*-EB1EGFP/EB3-mKate2 monopolar spindles using a widefield microscope. Upon blue light exposure (FITC channel 490 nm, 300 ms exposure), EB1 (purple) is acutely photoinactivated with EB3 (red; TRITC channel 555 nm) remaining to track microtubule ends. The TRITC channel was acquired before FITC thus providing a frame before photoinactivation (dark). Cells were imaged at 4 s intervals. Movies were analysed using TrackMate to detect EB3 comets (black circles) through time. A convex hull (yellow solid line) was applied on the detected spots to extract the monopolar spindle area. **B.** Representative widefield deconvolved single slice time-lapse images of H1299 *π*-EB1-EGFP/EB3-mKate2 cells undergoing whole-cell EB1 photoinactivation (blue border) at T = 0 s. Cells were treated with STLC 4 h before imaging to induce a monopolar state, and the indicated drugs just before imaging. **C.** Monopolar spindle area comparison before (0 min, grey) and after blue light exposure (10 min, blue) of H1299 *π*-EB1-EGFP/EB3mKate2 cells for the indicated conditions. Dark grey and blue dots indicate individual monopolar spindle area measurements with grey lines connecting data from the same cell. Black dots indicate outliers. Box plots show median along with interquartile range, while violin plots show the estimated kernel probability density. **D.** Line plot quantifying normalised monopolar spindle shrinkage exhibited by H1299 *π*-EB1-EGFP/EB3-mKate2 cells under blue light illumination with the indicated drugs. Shaded region displays 95 % confidence interval. **E.** Box plots quantifying the time taken to reach 85 % of the initial monopolar spindle area under EB1 photoinactivation (blue light) for the indicated conditions. Dots represent individual measurements from different cells. **F.** Box plots quantifying the time taken to reach 85 % of the initial detected EB3 comet number under EB1 photoinactivation (blue light) for the indicated conditions. Dots represent individual measurements from different cells. The analysis shown in this figure was performed after applying a moving average using three neighbouring values around the centre to smooth out area and comet change across time for each monopolar spindle. Sample size (n) indicated in (**C.**) is the same for (**D.**). Note that number of DMSO cells indicated in (**E.**) and (**F.**) is lower due to some cells not reaching 85 % by the end of acquisition. Statistical significance (*) or no significance (ns) in (**C.**) was determined through a paired t-test, in (**D.**) via a Mann-Whitney U test, in (**E.**) through a Kruskal-Wallis and a post-hoc Dunn’s test with Bonferroni correction, and in (**F.**) with a one-way ANOVA and post-hoc Tukey’s test. Data for Ciliobrevin-D 5 µM and Dynarrestin 2.5 µM are from two experimental repeats, while for other conditions data is from three repeats. T corresponds to the time at which the image was taken. Scale bar = 5 µm.

In contrast, monopolar spindle area and EB3 comet number showed a significant decrease after 10 min of EB1 photoinactivation in H1299 *π*-EB1-EGFP/EB3-mKate2 cells (Figure 3B), with the largest decrease observed with Dynarrestin at 25 µM (DMSO: 286.4 µm^2^ vs 249.2 µm^2^, Ciliobrevin-D 50 µM: 249.1 µm^2^ vs 179.0 µm^2^, Ciliobrevin-D 5 µM: 276.1 µm^2^ vs 187.3 µm^2^, Dynarrestin 25 µM: 251.2 µm^2^ vs 140.6 µm^2^, Dynarrestin 2.5 µM: 219.6 µm^2^ vs 148.2 µm^2^; median at 0 min vs 10 min) (Figure 3B and C). DMSO treatment findings align with superresolved images (Figure 1A), and spindle length reduction upon EB1 photoinactivation [51]. Importantly, dynein inhibition combined with EB1 photoinactivation led to a more severe reduction in monopolar spindle area defined by EB3 decorated microtubule-ends. Analysis through time showed a significant decrease (0.84-fold on average) at 10 min (Ciliobrevin-D 50 µM p = 0.026, Ciliobrevin-D 5 µM p = 0.033, Dynarrestin 25 µM p = 0.0079, Dynarrestin 2.5 µM p = 0.0071, vs DMSO; Mann-Whitney U test) (Figure 3D). The time to reduce to 85 % of the initial monopolar spindle area was significantly shorter for the higher concentration of Ciliobrevin-D (DMSO 514.3±193.7 s, Ciliobrevin-D 50 µM 303.7±228.3 s, Ciliobrevin-D 5 µM 365.5±159.5 s; mean±SD), and both concentrations of Dynarrestin (Dynarrestin 25 µM 324.8±254.6 s, Dynarrestin 2.5 µM 277.8±158.8 s; mean±SD) (Figure 3E). Similarly, the time required to reach 85 % of the initial EB3 comet number was significantly lower when dynein was inhibited (DMSO 547.3±192.0 s, Ciliobrevin-D 50 µM 361.3±272.0 s, Ciliobrevin-D 5 µM 399.0±160.5 s, Dynarrestin 25 µM 218.5±153.9 s, Dynarrestin 2.5 µM 264.9±102.8 s; mean±SD) (Figure 3F). As dynein inhibition alone does not significantly alter monopolar spindle area in the presence of EB1 (Figure S5D), we conclude that dynein contributes to EB1 independent astral microtubule growth.

### Dynein activity maintains astral microtubules independent of the canonical cortical dynein-NuMA-LGN pathway

We hypothesised that dynein’s role in maintaining astral microtubule length is independent of its cortical localisation as we observed asymmetrically positioned spindles, with one pole closer to the cortex displaying short aster, in the presence and absence of cortical dynein (Figure S4). To test this hypothesis, we removed cortical dynein in our EB1 photoinactivation studies by depleting LGN, the cortical platform for dynein [5, 8] (Figure 4A). Using LGN-RNAi suitable for live-imaging [9, 23], we first confirmed the effectiveness of LGN siRNA treatment in HeLa DHC-GFP cells, which allows direct visualisation of cortical dynein. LGN siRNA-treated (siLGN) HeLa DHC-GFP metaphase cells showed no cortical DHC-GFP signal (Figure 4B), indicating successful loss of cortical dynein.

**Figure 4.**
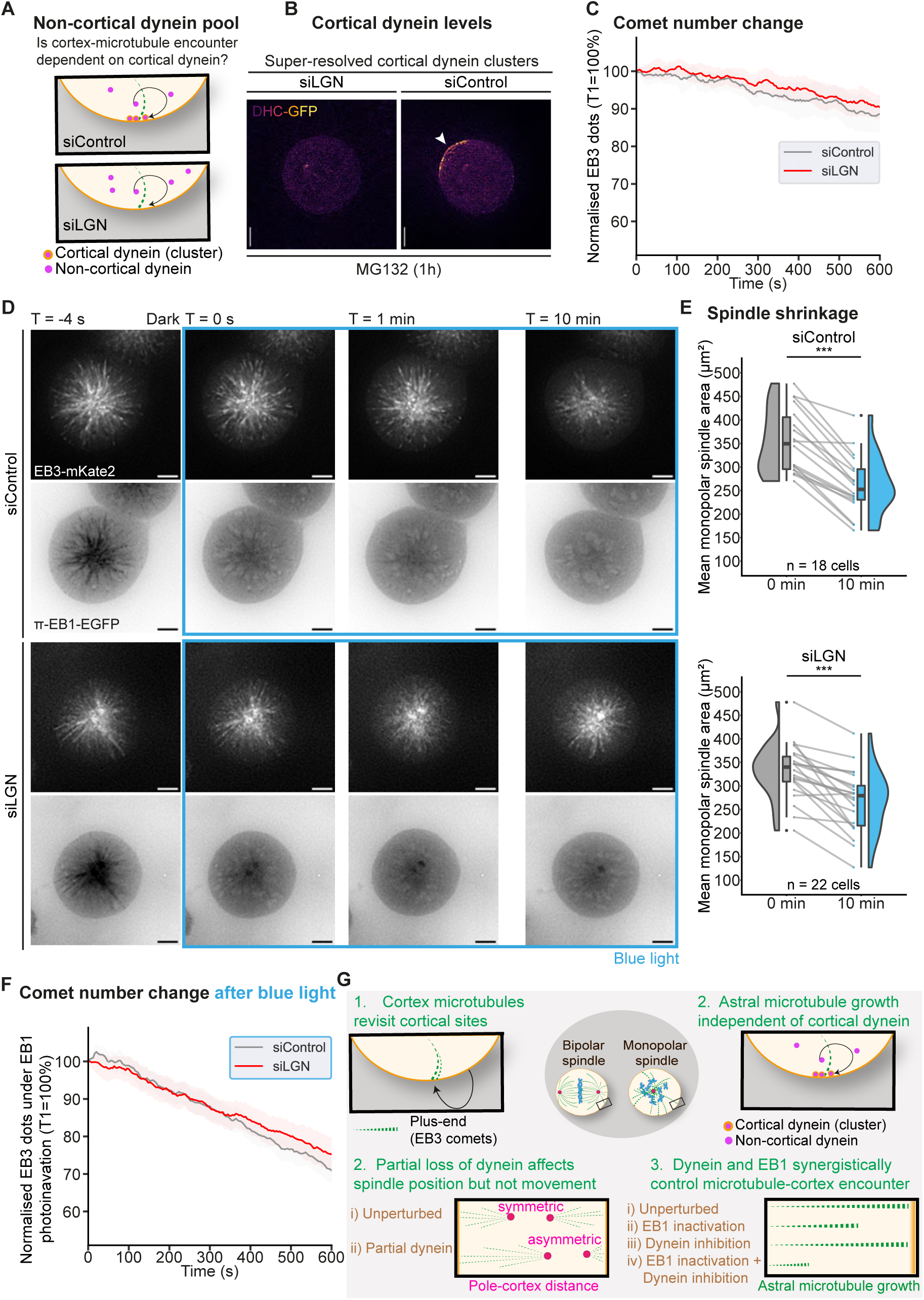
A role for noncortical dynein in astral microtubule maintenance. **A.** Schematic showing cortical and noncortical pools of Dynein in control but not LGN siRNA treated cells allowing the study of astral microtubule regulation by non-cortical dynein, in the absence of cortical dynein. **B.** Representative lattice SIM^2^ maximum projected images of metaphase-arrested HeLa DHC-GFP cells showcasing the depletion of cortical dynein (white arrowhead) following siLGN treatment. DHC-GFP intensities are used for pseudocolouring. DHC clusters at cortex are visible in control but not LGN siRNA treated cells. **C.** Line plot quantifying normalised change in EB3 comet number exhibited by H1299 *π*-EB1-EGFP/EB3mKate2 cells with the indicated siRNA treatments (n=13 Control siRNA and n=19 LGN siRNA treated cells). **D.**Representative widefield deconvolved single slice time-lapse images of H1299 *π*-EB1-EGFP/EB3-mKate2 cells undergoing whole-cell EB1 photoinactivation (blue border) at T = 0 s by being exposed to blue light (GFP channel, 470 nm). Cells were treated with STLC 4 h before imaging to induce a monopolar state, and the indicated siRNA. **B.** Monopolar spindle area comparison before (0 min, grey) and after blue light exposure (10 min, blue) of H1299 *π*-EB1-EGFP/EB3-mKate2 cells for the indicated conditions. Dark grey and blue dots indicate individual monopolar spindle area measurements with grey lines connecting data from the same cell. Box plots show median along with interquartile range, while violin plots show the estimated kernel probability density. Black dots indicate outliers. **F.** Line plot quantifying normalised detected EB3 comet number change exhibited by H1299 *π*-EB1-EGFP/EB3-mKate2 cells exposed to blue light (blue border) with the indicated siRNA treatments. Statistical significance (*) or no significance (ns) in (**E**) was determined through a paired t-test and in (**C**) and (**F**) via a Mann-Whitney U test. T corresponds to the time at which the image was taken relative to photoinactivation time. Scale bar = 5 µm. **G.** Illustrating how spindle positioning is influenced by microtubule-cortex encounter independent of cortical dynein. **1.** Growing astral microtubule-ends of monopolar and bipolar spindles retrace paths to specific cortical sites. **2.** Cortical dynein pulls at microtubules to move spindles towards the cortex, but surprisingly partial loss of dynein positions a pole closer to the cortex causing asymmetric spindle positioning (Red dots indicate spindle poles). **3.** This puzzle can be explained by dynein’s synergistic role with EB1 in maintaining microtubule growth which can regulate cortex encounter events of astral microtubule-ends (Green dashed lines represent growing microtubules). **4.** Maintaining astral microtubule growth is not reliant on the canonical cortex-bound pool of dynein. These explain how Dynein’s role in spindle pulling can be separated from its role in positioning that is linked to astral microtubule length maintenance. Thus, non-cortical and cortical dynein pools can be separately regulated to achieve asymmetric spindle positions to define cell division fates through non-overlapping developmental pathways.

In the absence of blue light, a small (12 %) but significant decrease in monopolar spindle area was observed at the end of 10 min movies in both conditions (siControl: 302.7±61.6 µm^2^ vs 265.7±73.9 µm^2^, siLGN: 325.9±66.2 µm^2^ vs 289.6±63.8 µm^2^; mean±SD at 0 min vs 10 min) (Figure S6A and S6B). Comparing the initial monopolar spindle area normalised to cell area showed a slight, non-significant decrease (8 %) following LGN siRNA-treatment (siControl 0.76±0.10, siLGN 0.70±0.09; mean±SD ratio) (Figure S6C), similar to observations upon dynein inhibition (Figure S5D). Both siControl and siLGN-treated monopolar spindles showed similar average area change across time (1.03-fold slope difference), with siLGN showing a slightly lesser area change (siControl 86.6 % vs siLGN 88.8 %) but not significantly different (p = 0.62; MannWhitney U test) (Figure S6D). The time to reach 85 % of initial monopolar spindle area was shorter for siControl (siControl 410 s vs siLGN 568 s; median), but not significantly different from siLGN (p = 0.091; unpaired t-test) (Figure S6E). Similar slight differences were observed in comet numbers, with siLGN showing a smaller but not significantly different change over time compared to siControl (Figure 4C and S6F). Thus in the presence of EB1, cells depleted of cortical dynein display no significant differences in microtubule growth compared to control cells.

In H1299 *π*-EB1-EGFP/EB3-mKate2 cells, monopolar spindle-to-cell area ratio showed no significant difference (p = 0.21; unpaired t-test) after LGN depletion (siControl 0.74±0.09, siLGN 0.71±0.10; mean±SD ratio) (Figure 4D and S7A). Following 10 minutes of EB1 photoinactivation, siControl spindles exhibited a slightly more severe decrease (28 % in median 0 min vs 10 min) compared to siLGN (18 % decrease in median 0 min vs 10 min) (Figure 4E). This trend was also reflected in the monopolar spindle area change over time, where siLGN cells showed a lesser but non-significant (p = 0.075; Mann-Whitney U test) reduction compared to siControl cells (1.09-fold difference) (Figure S7B). Time taken to reach 85 % of the initial monopolar spindle area or the initial number of EB3 comets after blue light exposure was not significantly different between siLGN and siControl cells (Figure S7D and S6F). In both cases siLGN cells showed lower area (siControl 324.7±156.6 s vs siLGN 457.5±252.1 s; mean±SD) and comet (siControl 338.7±159.7 s vs siLGN 381.8±240.7 s; mean±SD) reductions after EB1 photoinactivation (Figure S7C, S7D and 4F). Lack of accelerated monopolar spindle shrinkage after LGN-depletion is unlike observations after partial loss of dynein activity (Figure 3). Thus, non-cortical dynein synergises with EB1 to facilitate astral microtubule growth and encounter at the cortex.

## DISCUSSION

The spindle pulling role of the cortical force-generating machinery, dynein-dynactin with NuMALGN-G*α*_i_ complex, is well described [5–8, 10–12, 66]. We report a role for non-cortical dynein in facilitating astral microtubule-end growth to the cortex. We show that dynein and EB1 synergistically maintain astral microtubule growth, an event relevant to microtubule encounters at the cell cortex (Figure 4G). We show that this synergistic role for dynein with EB1 is distinct from its classical pulling role, using dynein inhibitor gradients and depleting the cortical dynein platform, LGN. The non-cortical role for dynein in maintaining astral microtubules sheds new light on how dynein ensures proper spindle positioning, beyond its classical role in spindle pulling.

A synergistic role for non-cortical dynein with EB1 aligns with *in vitro* studies. Purified dynein can stabilise plus-ends by exerting tension on protofilaments[27]. Nevertheless, barrier-bound high concentrations of dynein trigger microtubule catastrophe[7]. Thus, low-levels of non-cortical dynein can facilitate microtubule growth, search and encounter at cortex, where cortical clusters of NuMA-LGN-G*α*_i_[21] capture and pull at microtubules [57]. EB1 interacts with dynactin during prometaphase [67], a phase when microtubule interactions occur throughout the cortex to rotate spindles to a predetermined position[9, 59, 68]. The microtubule-cortex interaction facilitating role of EB1-dependent dynein, independent of EB3, is similar to other mitotic roles for EB1 known to be functionally independent of EB3 [44, 48, 50], highlighting the significance of the tip of the growing plus-end in the event.

Super-resolved microscopy movies show microtubule-ends retracing their path towards cortical sites in monopolar spindles, as in bipolar spindles (here and [4, 52, 53]). Disrupting cortex-microtubule interactions in different regions impact spindle movements differently, suggesting cortical dynein-dependent and independent roles for microtubule-ends in regulating spindle movements. This is consistent with cytoplasmic dynein driving oscillatory nuclear movements during prophase, as seen in fission yeast, where dynein localisation in astral microtubules can be visualised easily [69]. In small systems, spindle movements are more dependent on microtubule end-mediated push compared to large systems that rely on spindle pull[70]. In the pole closest to the cortex, we do not see a difference in spindle exit velocities or direction (Figure S3), ruling out spindle pulling or pushing forces in the asymmetric position seen after partial loss of dynein. We speculate that upon partial loss of dynein, poles remain proximal to the cortex at the end of a cortical dynein-mediated pull due to shorter asters from reduced astral microtubule growth. We favour reduced microtubule plus-end growth as the underpinning reason for asymmetric positioning as we find a significant reduction in EB3 comet numbers following partial dynein loss in EB1-photoinactivated monopolar spindles.

Dynein knock-out cells with turbulent spindles are reversed by coinhibiting Eg5 kinesin [71]. Here, partial dynein loss and Eg5 coinhibition exposes dynein’s synergistic action with EB1 in maintaining microtubule-end growth. In addition, partial loss of dynein separates the role of dynein in pole-cortex distance and symmetric positioning, from its known roles in spindle maintenance [71, 72] and displacement [9, 73]. Like dynein inhibition, MARK2/Par1 depletion can alter spindle positions without affecting spindle size or displacement rates: MARK2 depletion causes spindle off-centering along the equatorial axis [23], whereas dynein inhibition results in asymmetric spindle positioning (off-centering along the pole-to-pole axis). Moreover, the two treatments diverge in their effects on microtubule growth: MARK2 depletion causes microtubules to grow longer[23, 74], while dynein inhibition reduces microtubule length when combined with EB1 loss. Collectively, these observations highlight the close link between astral microtubule dynamics and spindle positioning. Uncovering dynein’s synergy with EB1, a protein hub with several partners mediating signalling and anchoring functions [25], highlights how non-cortical dynein and EB1 can integrate various cues from mechanical, polarity and developmental factors throughout the cell to correctly position the spindle.

## METHODS

### Cell culture

Cell cultures were maintained in sterile 60 mm Nunc™ cell culture dishes (10111351, Fisher Scientific). For larger cell cultures, 100 mm Corning (BC153, Appleton Woods) or 150 mm Nunc (15803947, Fisher Scientific) dishes were used. HeLa cell lines were cultured in Dulbecco’s Modified Eagle’s Medium (DMEM) (11500416, Fisher Scientific) supplemented with 10 % Fetal Bovine Serum (FBS) (10270106, ThermoFisher), 1 % Penicillin-Streptomycin (15140122, ThermoFisher), 1 % L-Glutamine (11539876, Fisher Scientific), and 0.1 % Amphotericin B (11510496, Fisher Scientific). H1299 cell lines were cultured in Roswell Park Memorial Institute (RPMI) 1640 Medium (11530586, Fisher Scientific) supplemented with 10 % FBS (10270106, ThermoFisher), 1 % Penicillin-Streptomycin (15140122, ThermoFisher), 1 % Minimum Essential Medium (MEM) Non-Essential Amino Acids Solution (11140050, ThermoFisher) and 0.1 % Amphotericin B (11510496, Fisher Scientific). Cell lines were cultured as a monolayer at 37 *^◦^*C and 5 % CO_2_.

### Cell line generation

For generating HeLa EB3-mKate2 and H1299 *π*-EB1-EGFP/EB3-mKate2 cell lines, unmanipulated HeLa and H1299 *π*-EB1-EGFP cells were seeded onto 60 mm dishes respectively. At 80 % confluency, each dish was transfected with 10 µg of plasmid encoding EB3-mKate2 cDNA using Dharmafect Duo Transfection Reagent (T2010-03, Horizon Discovery). Using Fluorescence-Activated Cell Sorting (FACS) with a BD FACSAria™ III Cell Sorter, cells with medium level of protein expression were collected in complete media and seeded in sterile 60 mm culture dishes. Stable cell pools were subsequently further isolated using antibiotic selection by supplementing complete culture media with G418 sulfate (108321, Sigma-Aldrich) at 0.8 mg/mL. Repeated rounds of FACS (at least 3, every 2-3 weeks), monitoring through a widefield microscope to assess the proportion of fluorescent cells with microtubule-end expression, and antibiotic selection (media supplemented with antibiotic replaced every 2-3 days) were carried out to ensure fluorescence stability.

### Live-cell Imaging

For live-cell deconvolution and super-resolution experiments, please see supplementary text.

### Transfection

siRNA transfection was carried out using Lipofectamine™ RNAiMAX Transfection Reagent (12323563, Fisher Scientific) according to the manufacturer’s protocol. Stealth RNAi™ siRNA Negative Control (10143902, Fisher Scientific) was used as depletion control. siRNA oligonucleotides dilutions were prepared in Opti-MEM, with a starting stock concentration of 10 µM. siRNA oligonucleotides were purchased from Sigma-Aldrich.

### Photoinactivation

In Super Resolution (SR) microsopy studies, illumination using 488 nm and 561 nm lasers was performed every 105 s with a ∼2 s gap for each frame (consisting of 13 phases for SIM). The blue light illumination block was subsequently triggered via the camera, acquiring a frame every 4 s with a 120 ms camera exposure (Figure 2A). For localised blue light illumination, in the SR regime, the 561 nm channel was imaged throughout the acquisition period, while a 473 nm laser was triggered to illuminate a specified ROI after a short delay (Figure 2B). In deconvolution microscopy studies, EB1 photoinactivation was achieved by blue light exposure triggered through the camera: for each frame the TRITC (555 nm) channel was captured first, followed by the FITC (490 nm) channel to induce EB1 photoinactivation (Figure 3A).

### Data visualisation and Statistics

Data was plotted using Python’s Seaborn or plotnine packages, Microsoft Excel, or GraphPad Prism 9. Statistical tests used are specified in the figure legends. The choice of statistical method was based on a pre-analysis of the underlying distribution of the data, such as through the Shapiro-Wilk test, ensuring that the assumptions of the tests were met. p-values are reported as follows: not significant (ns) for p > 0.05, (*) for p < 0.05, (**) for p < 0.01, (***) for p < 0.001.

## Data Availability

Original data sources will be made available through Zenodo and figshare.

## Acknowledgments

We thank Dr Torsten Wittman’s Lab for sharing their H1299 EGFP-Zdk1-EB1C (*π*-EB1) parent cell line. We thank members of the group of V.M.D. for useful discussions, Dr Gary Warnes (QMUL) for support with FACSORTing, and Dr Chengchen Wu (QMUL) for managing the superresolution microscopy facility. We acknowledge funding support from the Biotechnology and Biological Sciences Research Council (BBSRC) and InnovateUK to V.M.D. (BB/R01003X/1, BB/T017716/1, BB/W002698/1, and BB/X511067/1 and KTP012502), N.O. (BB/Y009002/1), C.E. (LIDo-iCASE studentship BB/T008709/1) and Regional England Investment Fund (2024) to V.M.D.

## Author Contributions

The project was conceptualised by VMD, and all experiments were designed together with CE. Experiments were conducted and analysed by CE, except in the case of figures 5F and 5G where data analysis method and investigation was conducted by NO. The original draft was prepared by CE with VMD’s support, and later versions were edited by VMD. VMD and CE discussed findings with NO and VMD group members. Figure panels were compiled by CE except for Figures 5E, 5H and 8 by VMD.

## Competing Interests

The authors declare no competing interests.

## Supplementary Methods

### Microscopy

Live-cell imaging of HeLa DHC-GFP and HeLa EB3-mKate2 cells was performed using a DeltaVision Core microscope (Applied Precision) which featured a dual-camera system comprised of a CoolSnap HQ2 CCD camera and a Cascade II EMCCD camera (Photometrics). Illumination was provided by a Xenon lamp, and different filter configurations (DAPI/FITC/TRITC/Cy5 for the CoolSnap HQ2 camera, GFP/mCherry for the Cascade II camera) were used accordingly. The imaging setup included an Olympus 100x/NA1.4 oil-immersion objective (UPLSAPO100XO). Z-stacks typically consisted of 3 Z-slices, with a spacing of 2 µm. Exposure times varied depending on the channel and camera used but were optimised to minimise both photobleaching and phototoxicity [75, 76]. Imaging was conducted in an incubation chamber maintaining the temperature at 37 *^◦^*C. For whole-cell photoinactivation experiments, cells were exposed to blue light (300 ms exposure) by acquiring the FITC channel (490 nm) or GFP channel (470 nm), following acquisition in the TRITC (555 nm) or mCherry (572 nm) channels, respectively. These time-lapse series were captured in a single focal plane with a 4 s interval, lasting for either 10 or 15 min. The pixel size for datasets captured with the Cascade II EMCCD camera was 0.128 20 µm, while for the CoolSnap HQ2 CCD camera it was 0.066 30 µm. The microscope was controlled through the softWorX software. Following acquisition, deconvolution was carried out using the enhanced ratio method for 10 cycles in the softWorX software.

Super-resolution live-cell imaging was performed using a structured illumination microscopy (SIM) system, Zeiss Elyra 7, equipped with a photokinetics module (Rapp OptoElectronic). The system included a 63x/NA1.4 oil immersion objective, a Duolink sCMOS camera adapter with two pco.edge 4.2 sCMOS cameras for simultaneous two-color acquisition, and a chamber incubation system maintaining temperature at 37 *^◦^*C. The setup featured a lattice SIM^2^ module that used a lattice spot illumination pattern instead of the traditional grid line SIM pattern, reducing phototoxicity and increasing imaging speed [77]. For whole-cell photoinactivation experiments, illumination was provided by 561 nm and 488 nm laser lines, paired with a secondary beam splitter long-pass filter (SBS LP560). Accordingly, blue and green channels were directed to one camera, while red and far-red channels went to the other. Acquisitions were performed on a single focal plane in illumination blocks. First block included both 561 nm and 488 nm channels (captured sequentially every frame) with a 2 s interval for three timepoints, followed by a longer 488 nm-only block, where frames were captured every 4 s. Thirteen phase images were captured for each frame with a 120 ms camera exposure time. For localised photoinactivation experiments, the photokinetics module was used to illuminate cells with a 473 nm laser while the 561 nm channel was acquired. The 473 nm laser intensity was adjusted using a neutral density filter, limiting transmission to 0.1 %. Laser power was set to 25 % for interphase cells and 5 % for mitotic cells. All lasers were calibrated according to the manufacturer’s guidelines, subsequently allowing laser illumination to be precisely localised to specified regions of interest (ROIs) created in Rapp OptoElectronic’s SysCon software. The photokinetics module was controlled through Rapp OptoElectronic’s SysCon software, which communicated with Zeiss’ Zen acquisition software via triggering. After acquisition, datasets were lattice SIM^2^ reconstructed with ‘Weak Live’ settings in Zen software [77]. The final pixel size after processing was 0.03 µm.

### Data Analysis

Manual quantifications, such as spindle displacement in 2D, spindle pole-pole distance, spindle-pole cortex distance, and cell area measurements, were performed using Fiji [78]. Additional image processing, including maximum Z-projections, temporal color projections, channel LUT specification, brightness and contrast adjustments, and exporting, were also done using Fiji.

Semi-automatic tools tested for measuring monopolar spindle areas included Cellpose [79], ilastik [80], u-track [81, 82], TrackMate [65, 83], and conventional intensity thresholding. For Cellpose, the ‘cyto3’ model [84] was tested via a Google Colab notebook [85], with the diameter parameter adjusted according to the input image. For ilastik, the ‘Pixel Classification’ workflow was used by following the developers’ documentation [86]. In u-track, the ‘Comet detection’ workflow was applied with these optimised parameters; Band-pass filter parameters: Low-pass Gaussian standard deviation = 1, High-pass Gaussian standard deviation = 3; Watershed segmentation parameters: Minimum threshold = 2.65, Threshold step size = 1. For TrackMate, several spot detectors were tested: 1) Difference of Gaussian (DoG), which makes it easier to detect structures like spots or edges by subtracting a more blurred version (higher *σ*) of an image from a less blurred version (lower *σ*), highlighting areas where the intensity changes rapidly [87, 88] (DoG is also used in u-track’s first band-pass filter step); 2) Laplacian of Gaussian (LoG), which first applies a Gaussian blur to smooth the image followed by the Laplacian operator (a second-order differential that calculates the difference between a pixel’s intensity and the average intensity of its neighbours) [89, 90] to identify regions of sharp intensity changes or local maxima [91, 92]; and 3) Hessian detector which utilises the determinant of the Hessian matrix to consider intensity changes in all directions around a pixel thereby providing a measure of the local curvature in the image [93, 94], subsequently identifying blobs and edges [95–97]. For all TrackMate spot detectors, the estimated object diameter was set to 1 µm, and the Quality metric was adjusted accordingly. In the conventional intensity thresholding tests, TrackMate’s Thresholding detector was used, with parameters fine-tuned based on the input image.

After testing, LoG’s detector was selected for analysing changes in monopolar spindle areas. In cases where the SNR was low, ROIs and X/Y filters were defined to exclude false-positive background spots. The Quality metric was fine-tuned for each cell individually. Once TrackMate completed the detection workflow, masks containing the detected spots as single pixels were exported. To calculate the monopolar spindle area, a custom Groovy script was used in Fiji. This script applied a 2D convex hull algorithm to generate the smallest convex polygon enclosing all detected binary spots. The script features a GUI that prompts the user to specify input and output folder paths, and generates a CSV file with area measurements and the X/Y centroid coordinates of the convex hull-drawn polygon.

## Supplemental Figures

**Figure S1.**
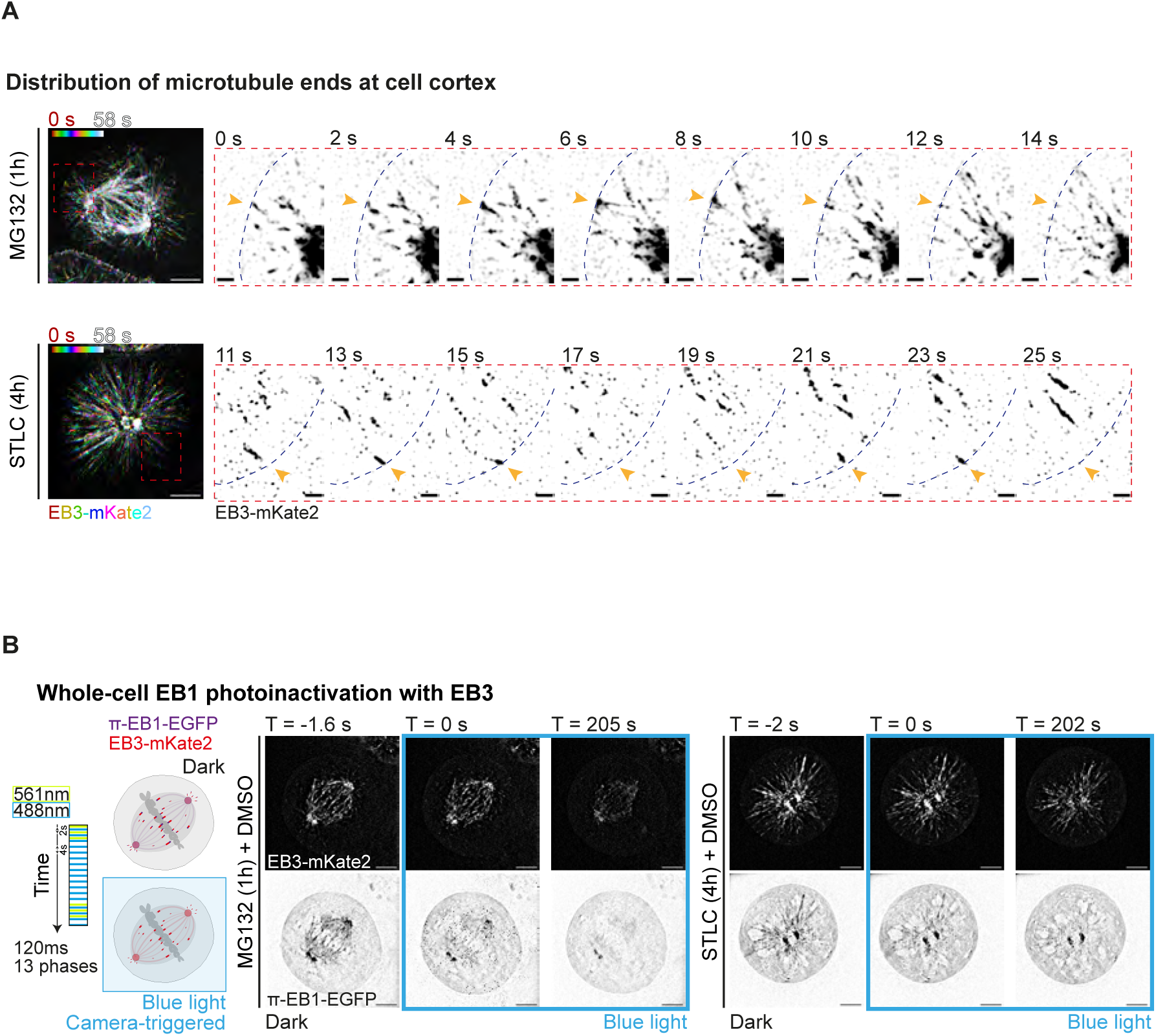
EB3 comets retrace paths to cortical sites in bipolar and monopolar spindles. **A** Representative temporal-colour coded lattice SIM^2^ images of an H1299 *π*EB1-EGFP/EB3-mKate2 bipolar (treated with MG132 for 1 h before imaging and 50 nM SiRTubulin dye just before imaging) and monopolar (treated with STLC for 4 h before imaging) spindle. Cells were imaged for 13 phases every 2 s for a total time of 58 s, capturing only the 561 nm (EB3-mKate2) channel. Red dotted boxes are zoomed time-lapse sequences (inverted contrast) of the regions indicated, showing microtubule ends reaching the same area (orange arrowhead) on the cortex (blue dotted line). Data representative of 16 bipolar and 10 monopolar spindles across two experimental repeats. Scale bar = 5 µm, 1 µm for zoomed images. **B.** Schematic illustrating camera-triggered whole cell EB1 photoinactivation (blue border) in H1299 *π*-EB1-EGFP/EB3-mKate2 cells. Upon exposure to blue light (488 nm), EB1 (purple) is acutely photoinactivated, and EB3 (red) remains transiently to track microtubule-ends. Next to the schematic are representative lattice SIM^2^ single slice time-lapse images of H1299 *π*-EB1EGFP/EB3-mKate2 cells undergoing whole-cell EB1 photoinactivation (blue border) at T = 0 s. Cells were treated with MG132 1 h before imaging to arrest cells in metaphase, or STLC 4 h before imaging to induce a monopolar state. Bipolar spindle images are representative of 32 cells across three repeats. Monopolar spindle images are representative of 15 cells across two repeats.

**Figure S2.**
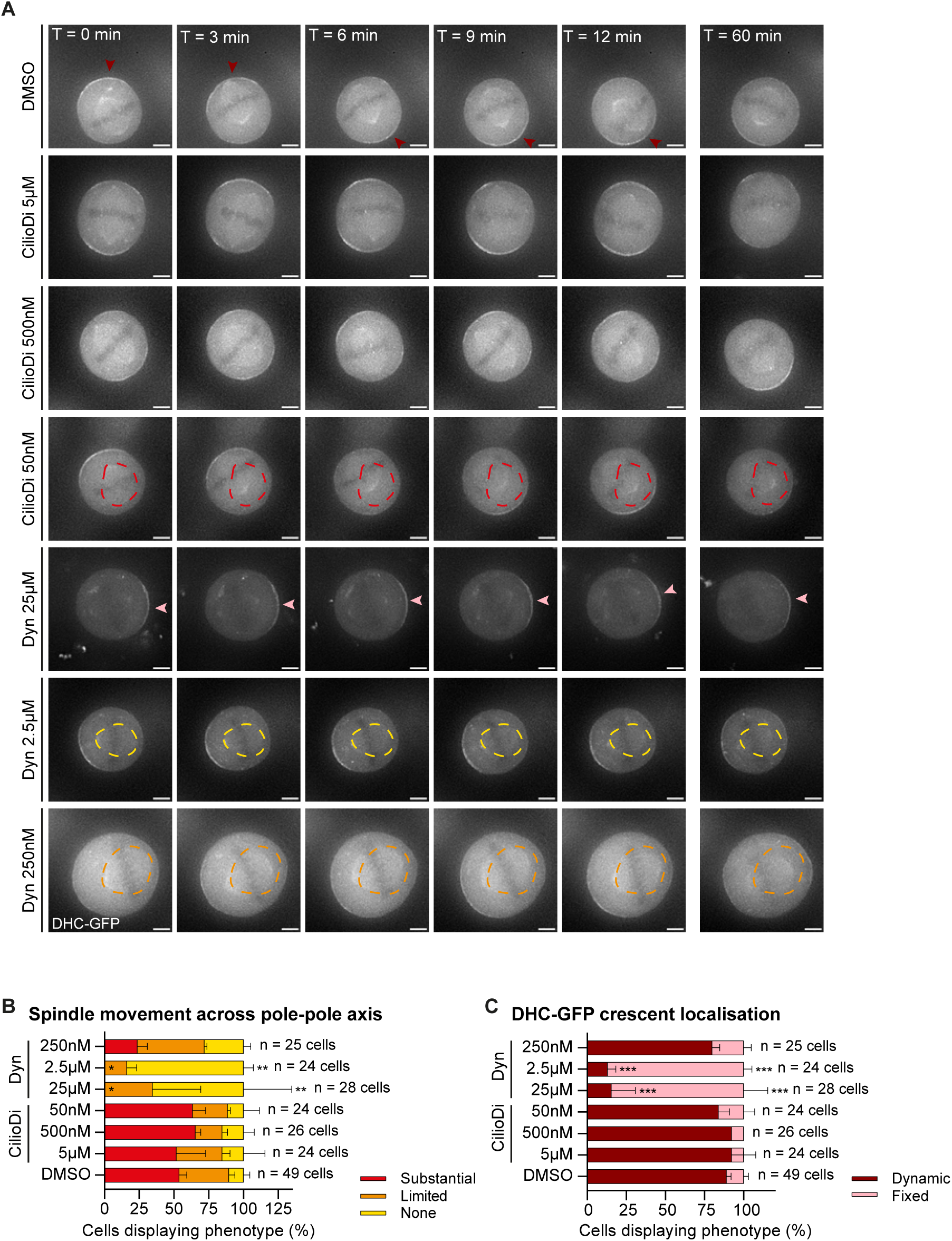
Dynein inhibitor gradient separates dynein’s roles in spindle movements and spindle positioning **A.** Representative widefield deconvolved maximum projected time-lapse images of HeLa DHC-GFP cells from each condition. Dark red arrowheads indicate a dynamic DHC-GFP cortical crescent, while light red arrowheads indicate a fixed DHC-GFP cortical crescent. Red, orange and yellow dotted lines trace the spindle’s position at the start of acquisition (T = 0 min), showcasing spindles exhibiting substantial (more than 2 spindle oscillations across the pole-pole axis), limited (at least 1 spindle oscillation) and no (None) movement throughout 60 min respectively. **B.** Stacked bar chart quantifying the proportion of spindle movements across the spindle pole-pole axis seen in cells treated as indicated. **C.** Stacked bar chart quantifying proportion of cortical DHC crescent localisation seen in cells treated as indicated. In (**B**) and (**C**) statistical significance (*) was determined by one-way ANOVA and post-hoc Tukey’s test. Statistical comparisons were only against DMSO. Normality was confirmed with a Shapiro-Wilk test for both (**B**) and (**C**). T corresponds to the time at which the image was taken. Scale bar = 5 µm. Sample size (n) as indicated. Data is presented as mean±s.e.m. across two experimental repeats.

**Figure S3.**
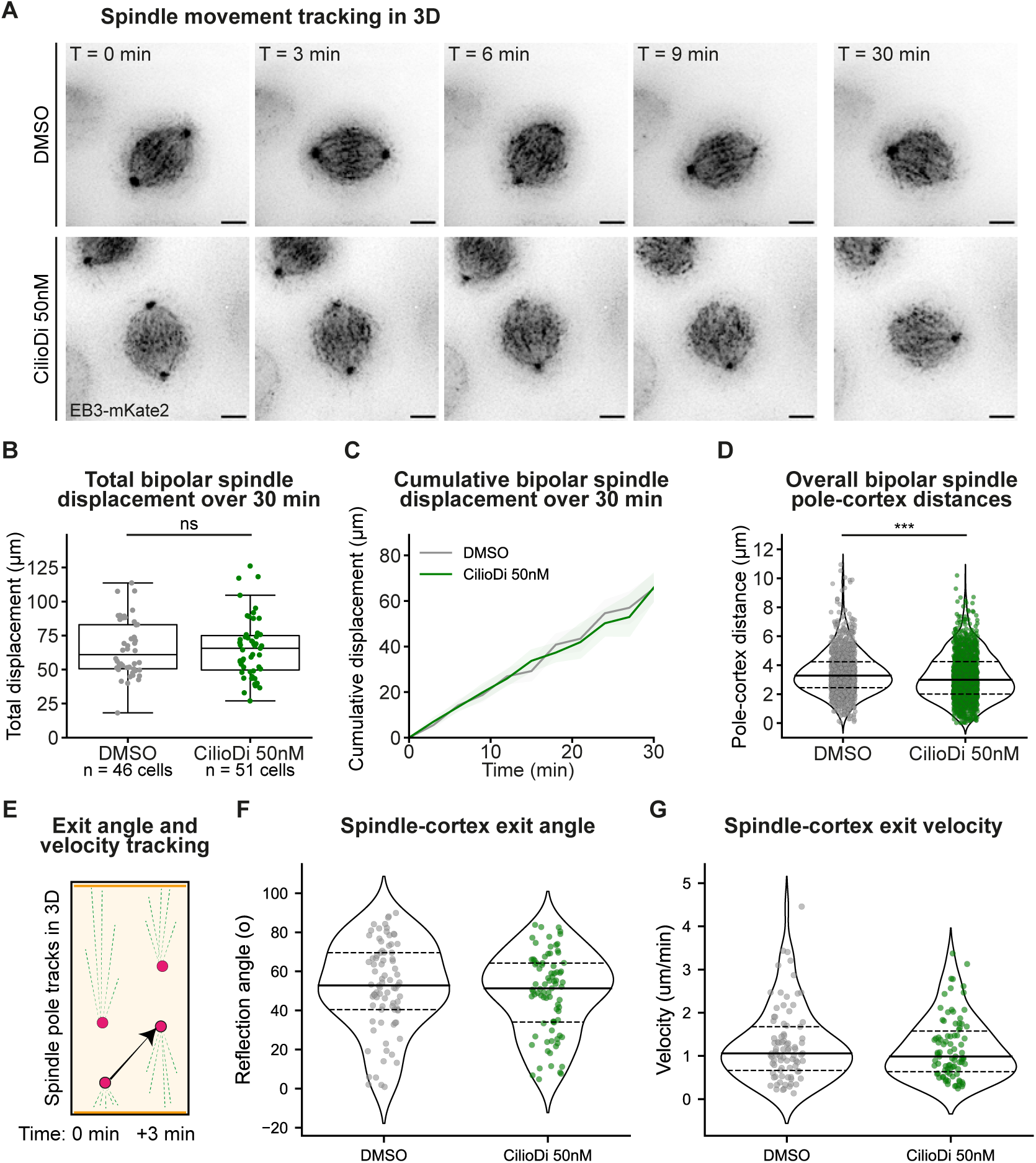
Ciliobrevin-D treatment causes reduced pole-to-cortex distance without altering total spindle displacement **A.** Representative deconvolved maximum projected time-lapse images of HeLa EB3-mKate2 cells treated with MG132 1 h before imaging to arrest cells in metaphase, and either DMSO or Ciliobrevin-D 50 nM just before imaging. **B.** Box plot quantifying the total displacement showed by bipolar spindles treated with DMSO and Ciliobrevin-D 50 nM over a 30 min imaging period. Dots indicate individual measurements. **C.** Line plots quantifying the average cumulative displacement exhibited by bipolar spindles treated with DMSO and Ciliobrevin-D 50 nM. Shaded region displays 95 % confidence interval. **D.** Violin plots showing the distribution of pole-cortex distances in DMSO and Ciliobrevin-D 50 nM-treated bipolar spindles. Dots represent measurements from both spindle poles across all timepoints. Dotted lines indicate the quartiles, while the solid lines show the median. **E** Illustration of spindle pole track measurements to study the behaviour of asymmetrically positioned bipolar spindles. Spindle poles (red) closest to the cortex (orange) were tracked for the rate and angle in which they moved away from cortex. 3D data on spindle pole positions was acquired using SpinX software outputs of spindle tracks from time-lapse images acquired once every three minutes. **F and G** Violin plot showing the distribution of angles (F) and velocities (G) of spindle poles moving away from the cell cortex (n=48 DMSO-treated and n=52 Ciliobrevin treated cells). Angles and velocities were measured between the spindle axis and the direction of the spindle reflection away from the cortex within two subsequent time frames (3 minutes). **H** Schematic summarising the impact of partial dynein inhibition and acute EB1 inactivation on spindle movements and astral microtubules in bipolar and monopolar spindles. (i) Partial inhibition of dynein reduces spindle pole-to-cortex distances without altering canonical dynein function in cortical pulling associated spindle displacement or velocities. (ii) EB1 photoinactivation reduces astral microtubule despite the presence of EB3. (iii) EB1 inactivation and Dynein inhibition synergistically reduce astral microtubule length and microtubule reach to the cell cortex.

**Figure S4.**
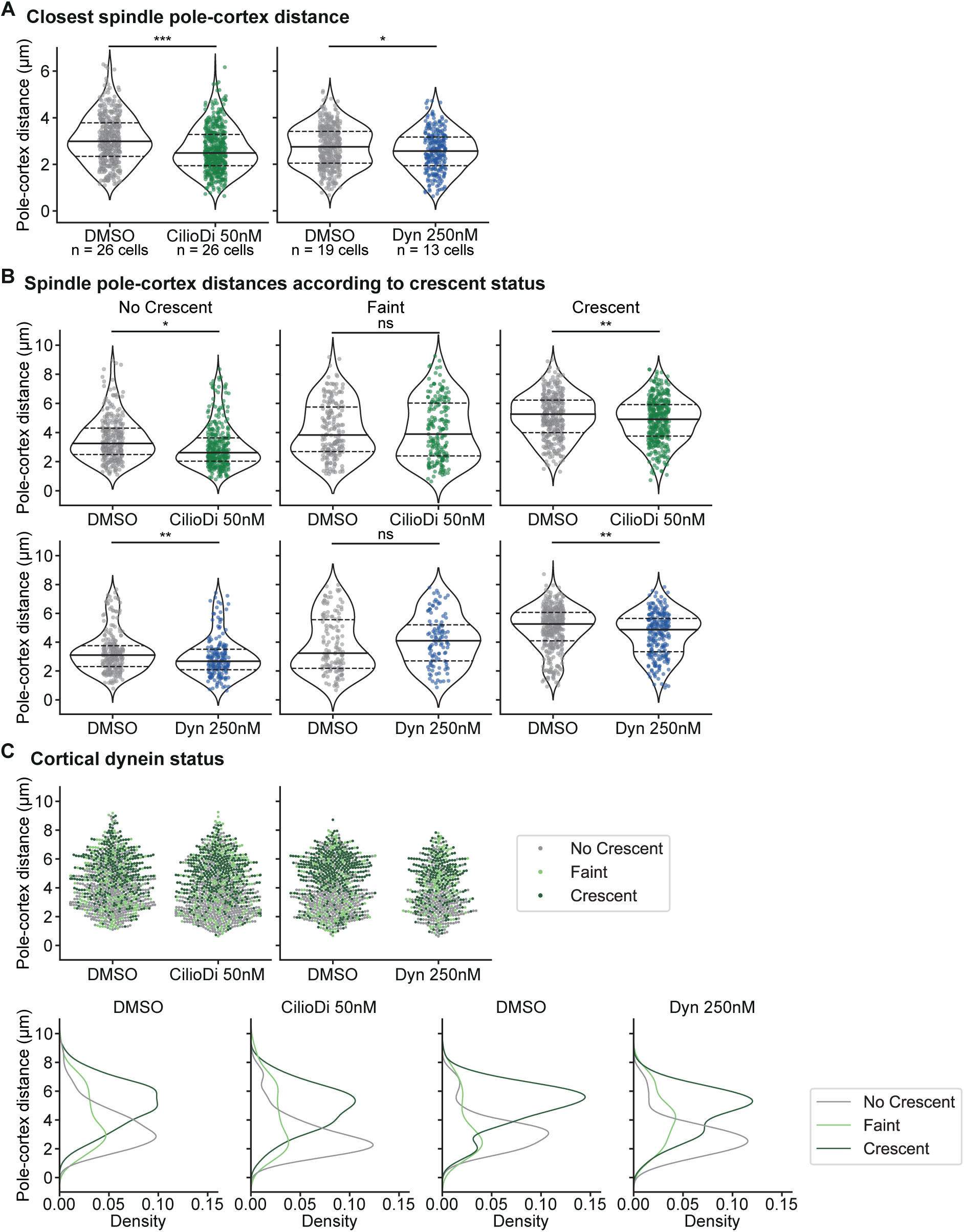
Partial inhibition of dynein results in reduced spindle pole-cortex distances with no changes to cortical DHC-GFP localisation. A. Violin plots showing the distribution of pole-cortex distances closest to the cortex in HeLa DHC-GFP DMSO, Ciliobrevin-D 50 nM and Dynarrestin 250 nM-treated spindles. Dots represent measurements from the spindle pole closest to the cortex at each timepoint. Dotted lines indicate the quartiles, while the solid lines show the median. B. Violin plots displaying the distribution of pole-cortex distances closest to the cortex in DMSO, Ciliobrevin-D 50 nM and Dynarrestin 250 nM-treated spindles. Dots represent measurements from the spindle pole closest to the cortex at each timepoint. C. Top: Swarm plots illustrating the distribution of spindle pole-cortex distances relative to DHC crescent status in spindles treated with DMSO, Ciliobrevin-D 50 nM and Dynarrestin 250 nM. Bottom: Corresponding KDE curve plots showing the probability density of spindle pole-cortex distances for each crescent status in each condition. Statistical significance (*) in (A) was determined by a Mann-Whitney U test. In (B) statistical significance and no statistical significance (ns) was determined either through an unpaired t-test after confirming equal variances with Levene’s test, or a Mann-Whitney U test. Normality was checked with a Shapiro-Wilk test. Sample size (n) as displayed in (A) is the same for (B) and (C), across 3 experimental repeats.

**Figure S5.**
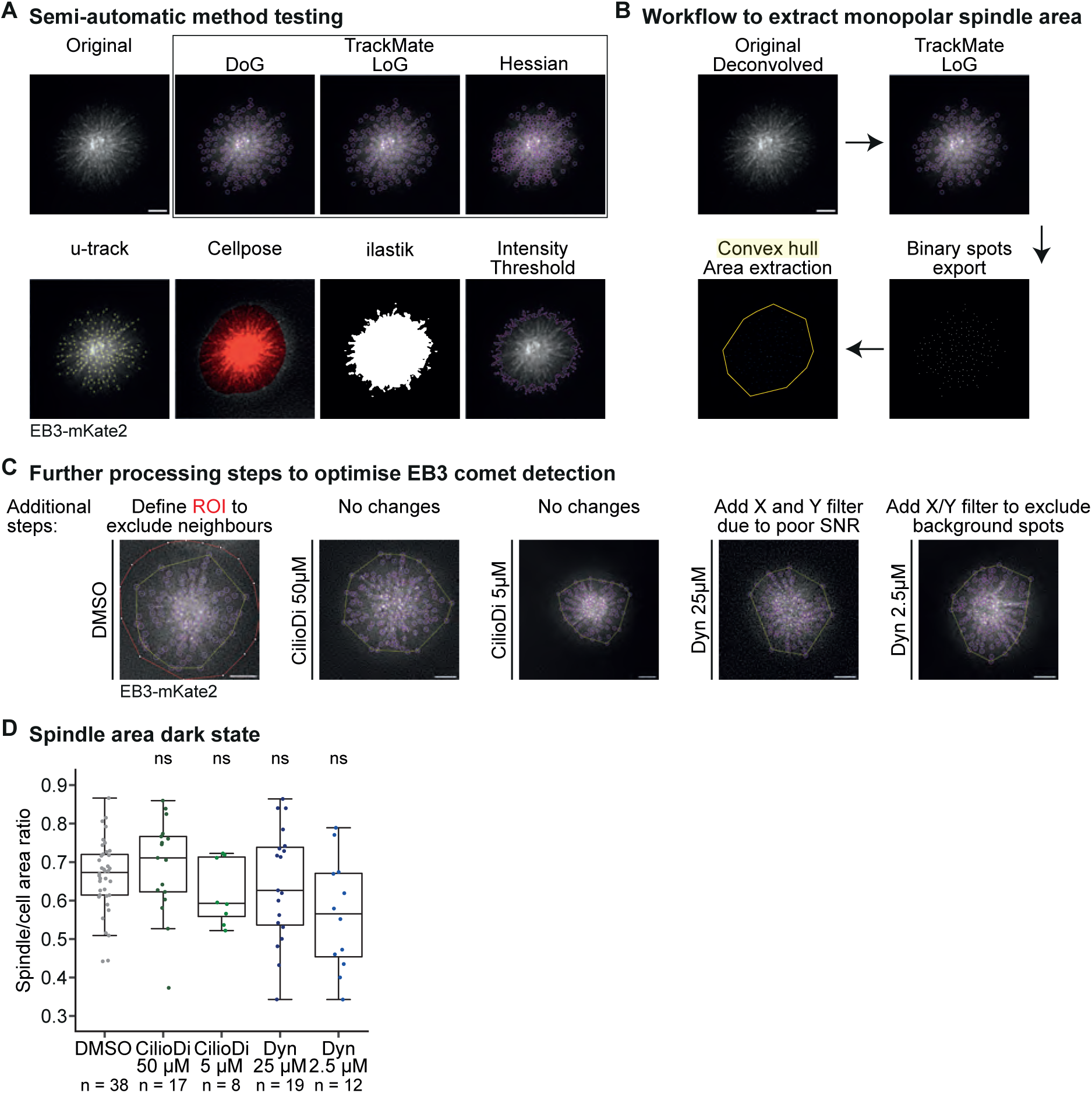
Dynein inhibition in the presence of EB1 does not significantly alter the length of astral microtubules reaching the cortex. **A.** Representative widefield deconvolved single-slice images of an H1299 *π*-EB1-EGFP/EB3-mKate2 cell demonstrate various methods tested for detecting spindle area in a monopolar spindle. The semi-automatic methods included different spot detectors provided by TrackMate (Difference of Gaussian, DoG; Laplacian of Gaussian, LoG; Determinant of Hessian, Hessian; purple circles), u-track (yellow circles), Cellpose (red area), illastik (white pixels) and conventional intensity thresholding (purple outline). **B.** Schematic depicts workflow to extract monopolar spindle area from the EB3-mKate2 channel. This includes 1) using TrackMate’s LoG detector with an estimated object diameter of 1 µm, a median filter, sub-pixel localisation and a cell-specific quality metric threshold (based on image background) on deconvolved H1299 EB3-mKate2 channel images; 2) exporting the detected spots (purple circles) as a binary mask (white pixels); and 3) applying a convex hull via a Fiji script to extract area (yellow solid line) and centroid XY coordinates. **C.** Representative images from H1299 *π*-EB1-EGFP/EB3-mKate2 monopolar spindles (induced through STLC treatment for 4 h before imaging) treated with either DMSO (Control), or Ciliobrevin-D 50 µM and 5 µM, or Dynarrestin 25 µM and 2.5 µM just before imaging. Images show additional processing steps to improve EB3 comet detection (purple circles) through TrackMate, and in turn the area measurement (yellow solid lines). These included defining an ROI (red line with points) to exclude neighbouring cells and adding X/Y coordinate filters to remove any background spots detected due to poor SNR or artefacts. **D.** Box plot quantifying initial monopolar spindle/cell area ratio for the conditions indicated. Dots represent individual measurements. No significance (ns) was determined using one-way ANOVA after confirming normality with Shapiro-Wilk test. Number of cells (n) as indicated. Ciliobrevin-D 5 µM and Dynarrestin 2.5 µM are across two experimental repeats while rest of conditions are across three. Scale bar = 5 µm.

**Figure S6.**
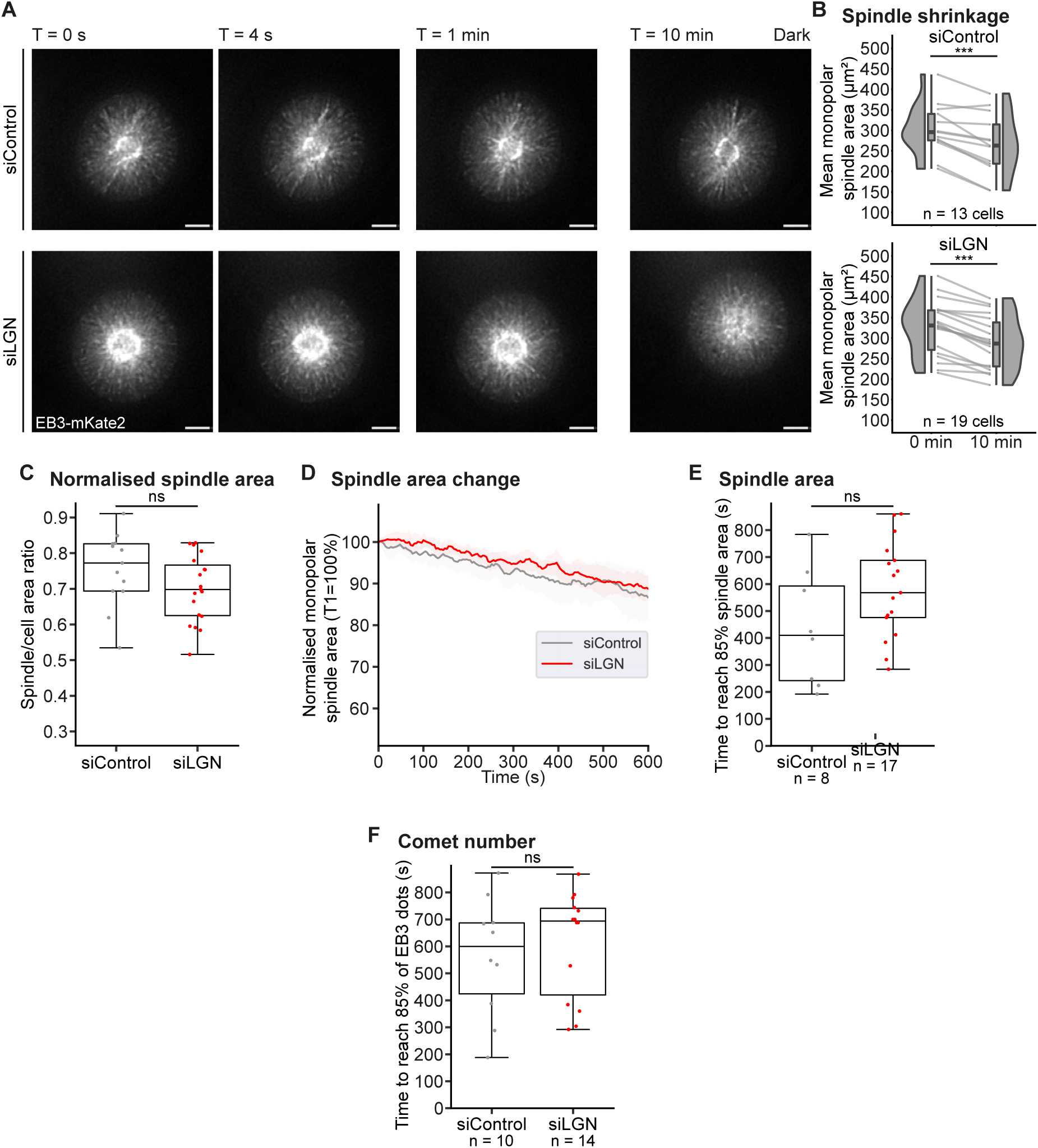
Control and LGN-depleted monopolar spindles behave similarly in the presence of EB1. **A**. Representative widefield deconvolved single slice time-lapse images of H1299 *π*-EB1EGFP/EB3-mKate2 cells treated with STLC 4 h before imaging and the indicated siRNA. Only EB3-mKate2 (mCherry channel, 572 nm) and cell cortex (DIC channel) were captured. **B.** Monopolar spindle area comparison at 0 min and 10 min of H1299 *π*-EB1-EGFP/EB3-mKate2 cells for the indicated conditions. Dots indicate individual monopolar spindle area measurements with grey lines connecting data from the same cell. Box plots show median along with interquartile range, while violin plots show the estimated kernel probability density. **C.** Box plot quantifying initial monopolar spindle/cell area ratio for the conditions indicated. Dots represent individual measurements. **D.** Line plot quantifying normalised monopolar spindle area change exhibited by H1299 *π*-EB1-EGFP/EB3-mKate2 cells with the indicated siRNA treatments. Shaded region displays 95 % confidence interval. **E.** Box plots quantifying the time taken to reach 85 % of the initial monopolar spindle area for the indicated conditions. Dots represent individual measurements from different cells. **F.** Box plots quantifying the time taken to reach 85 % of the initial detected EB3 comet number for the indicated conditions. Dots represent individual measurements from different cells. The analysis shown in this figure was performed after applying a moving average using three neighbouring values around the centre to smooth out area and comet change across time for each monopolar spindle. Sample size (n) indicated in (**B**) is the same for (**C**) and (**D**). Note that number of cells indicated in (**E**) and (**F**) is lower due to some cells not reaching 85 % by the end of acquisition. Statistical significance (*) or no significance (ns) in (**B**) was determined through a paired t-test, in (**C**) and (**E**) through an unpaired t-test, and in (**D**) and (**F**) via a Mann-Whitney U test. Data is across three experimental repeats. T corresponds to the time at which the image was taken. Scale bar = 5 µm.

**Figure S7.**
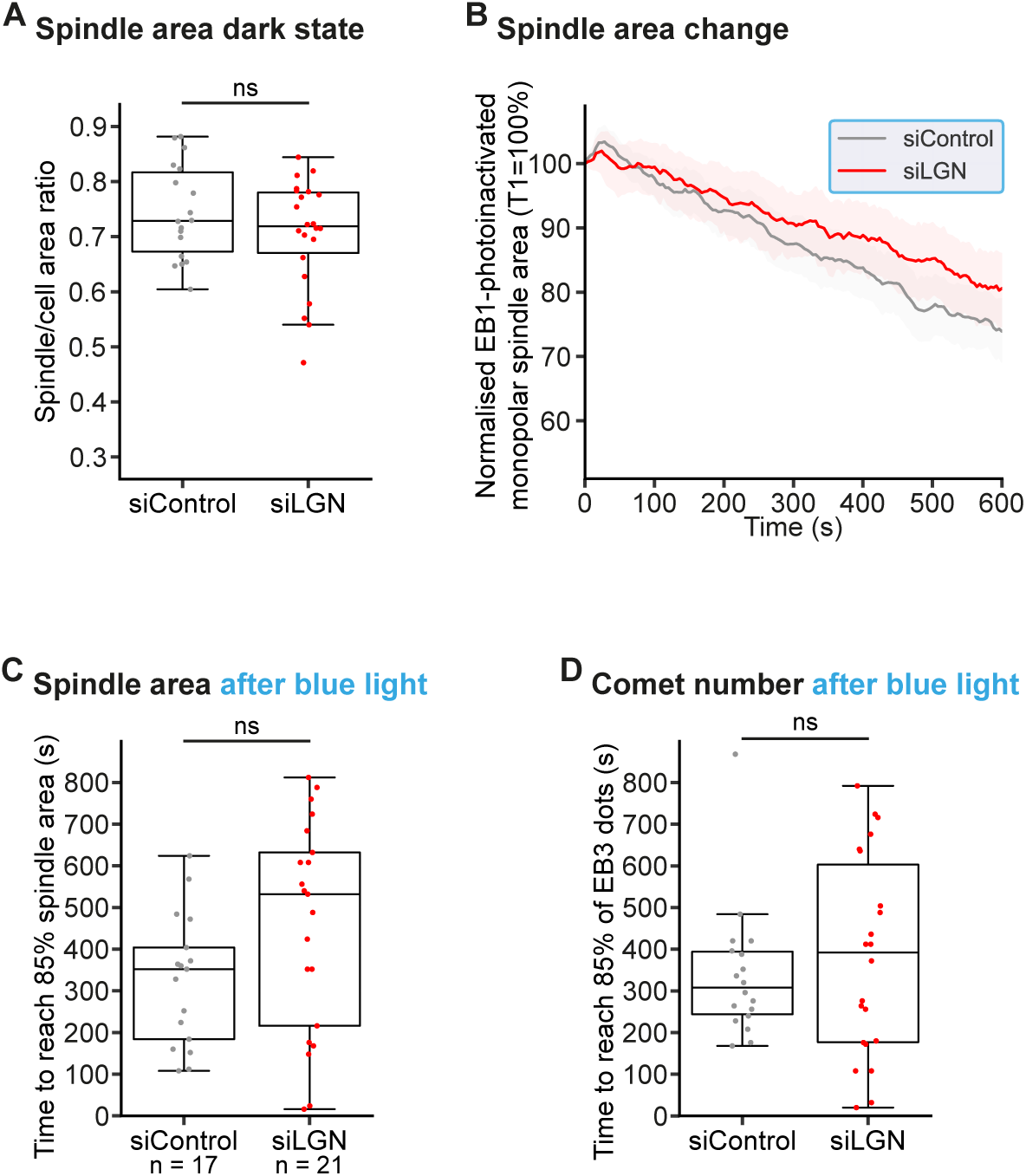
Monopolar spindles depleted of LGN resistant astral microtubule shrinkage upon EB1 photoinactivation. **A.** Box plot quantifying initial monopolar spindle/cell area ratio for the conditions indicated. Dots represent individual measurements. **B.** Line plot quantifying normalised monopolar spindle area change exhibited by H1299 *π*-EB1EGFP/EB3-mKate2 cells undergoing EB1 photoinactivation (blue border), with the indicated siRNA treatments. Shaded region displays 95 % confidence interval. **C.** Box plots quantifying the time to reach 85 % of the initial monopolar spindle area following EB1 photoinactivation for the indicated conditions. Dots represent individual measurements from different cells. **D.** Box plots quantifying the time taken to reach 85 % of the initial detected EB3 comet number after EB1 photoinactivation for the indicated conditions. Dots represent individual measurements from different cells. The analysis shown in this figure was performed after applying a moving average using three neighbouring values around the centre to smooth out area and comet change across time for each monopolar spindle. Sample sizes (n) for (**A**), (**B**) and (**D**)are as in (**Figure 4E**). Note that the number of cells indicated in (**C**) is lower due to some cells not reaching 85 % by the end of acquisition. Statistical significance (*) or no significance (ns) in (**A**) and (**C**) was determined through a unpaired t-test and in (**B**) and (**D**) via a Mann-Whitney U test. Data is across three experimental repeats. T corresponds to the time at which the image was taken. Scale bar = 5 µm.

## Notes

### Competing Interest Statement

The authors have declared no competing interest.

